# Variability in the phytoplankton response to upwelling across an iron limitation mosaic within the California Current System

**DOI:** 10.1101/2023.09.22.558967

**Authors:** YuanYu Lin, Olivia Torano, Logan Whitehouse, Emily Pierce, Claire P. Till, Matthew Hurst, Robert Freiberger, Travis Mellett, Maria T. Maldonado, Jian Guo, Mariam Sutton, David Zeitz, Adrian Marchetti

## Abstract

Coastal upwelling currents such as the California Current System (CCS) comprise some of the most productive biological systems on the planet. Diatoms, a distinct taxon of phytoplankton, dominate these upwelling events in part due to their rapid response to nutrient entrainment. In this region, they may also be limited by the micronutrient iron (Fe), an important trace element primarily involved in photosynthesis and nitrogen assimilation. The mechanisms behind how diatoms physiologically acclimate to the different stages of the upwelling conveyor belt cycle with respect to Fe limitation remains largely uncharacterized. Here, we explore their physiological and metatranscriptomic response to the upwelling cycle with respect to the Fe limitation mosaic that exists in the CCS. Subsurface, natural plankton assemblages that would potentially seed surface blooms were examined over wide and narrow shelf regions. The initial biomass and physiological state of the phytoplankton community had a large impact on the overall response to simulated upwelling. Following on-deck incubation under varying Fe physiological states, our results suggest that diatoms quickly dominated the blooms by “frontloading” nitrogen assimilation genes prior to upwelling. However, diatoms subjected to induced Fe limitation exhibited reductions in carbon and nitrogen uptake and decreasing biomass accumulation. Simultaneously, they exhibited a distinct gene expression response which included increased expression of Fe-starvation induced proteins and decreased expression of nitrogen assimilation and photosynthesis genes. These findings may have significant implications for upwelling events in future oceans, where changes in ocean conditions are projected to amplify the gradient of Fe limitation in coastal upwelling regions.

## Introduction

Eastern Boundary Upwelling Currents (EBUC), while comparatively smaller in size to other regions of the ocean, are disproportionately some of the most productive ecosystems on the planet [Capone and Hutchins 2013, Carr 2002]. Characterized by equatorward wind patterns that result in a net offshore transport of surface water and the upwelling of cold nutrient-rich water from below [Huyer 1983], these regions provide for extensive phytoplankton blooms that not only support the local food web but also mediate the flux of carbon into the deep ocean [Capone and Hutchins 2013]. The California Current System (CCS), one such EBUC, is particular in its irregular bathymetry and seasonality, where the frequency of upwelling is enhanced more often during the spring and summer [Closset et al. 2021]. The unusual qualities of the CCS are furthermore pronounced by its Fe limitation mosaic [Bruland et al. 2001, Hutchins et al. 1998], in which regions with wide and shallow continental shelves are mostly considered Fe-replete, while those with narrow and deeper continental shelves have been found to experience Fe limitation of phytoplankton growth [Bruland et al. 2001]. This is explained by the fact that wide and shallow continental shelves act as Fe traps [Capone and Hutchins 2013]. During the upwelling season, Fe-rich sediments from the bottom are more likely entrained into surface waters to seed phytoplankton blooms, while there is an inadequate supply of Fe to complement high macronutrient concentrations in narrow shelf regions because upwelling is focused off the shelf and over the deeper continental slope [Bruland et al. 2001, Capone and Hutchins 2013]. Occasionally when these trends aren’t observed, it is usually due to mesoscale features such as eddies moving water that was recently upwelled inland of the shelf offshore, or vice versa [Till et al. 2019].

Within this complex system lies a sequence of idealized light and nutrient zones (Fig. 1). On the surface, upwelled phytoplankton physiologically acclimate to optimal light and nutrient conditions (high light, high nutrients) and accelerate nitrogen rate processes relative to carbon rate processes, leading to enhanced macromolecule synthesis and growth referred to as ‘shift-up’ [Wilkerson and Dugdale 1987]. The increased algal growth processes exhibited during this stage is matched by the increase of their biomass and the formation of phytoplankton blooms. By maintaining high transcript abundances of nitrogen assimilation genes even in the seed populations, diatoms can ensure a more rapid physiological response to upwelling compared to other phytoplankton taxa. This advantageous “frontloading” of nitrogen assimilation machinery allows them to rapidly dominate these blooms in coastal upwelling areas [Lampe et al. 2018], and as noted by Margalef’s Mandala principle, exhibit high specific growth rates and thrive under high turbulence (upwelling) [Margalef 1978]. A course of succession follows as other phytoplankton taxonomic groups exhibit a slower reaction and response to the upwelling. As nutrient concentrations are rapidly depleted (high light, low nutrients), the cells undergo a ‘shift-down’ response, in which rate processes decrease further downstream. Phytoplankton will sink throughout this cycle, and may be re-upwelled in the next upwelling event [Wilkerson and Dugdale 1987]. Naturally, phytoplankton seed populations are established from the sinking communities as they are advected offshore. During this stage, the aging waters may be increasingly nitrate limited, and cells increase their C:N ratios above that of the Redfield ratio (6.6:1) as they sink below the euphotic zone. Once at depth (low light, high nutrients), diatoms are set apart by the differential expression of nitrogen transporters and other nitrogen assimilation genes compared to other phytoplankton taxa [Lampe et al. 2021].

**Figure 1.**
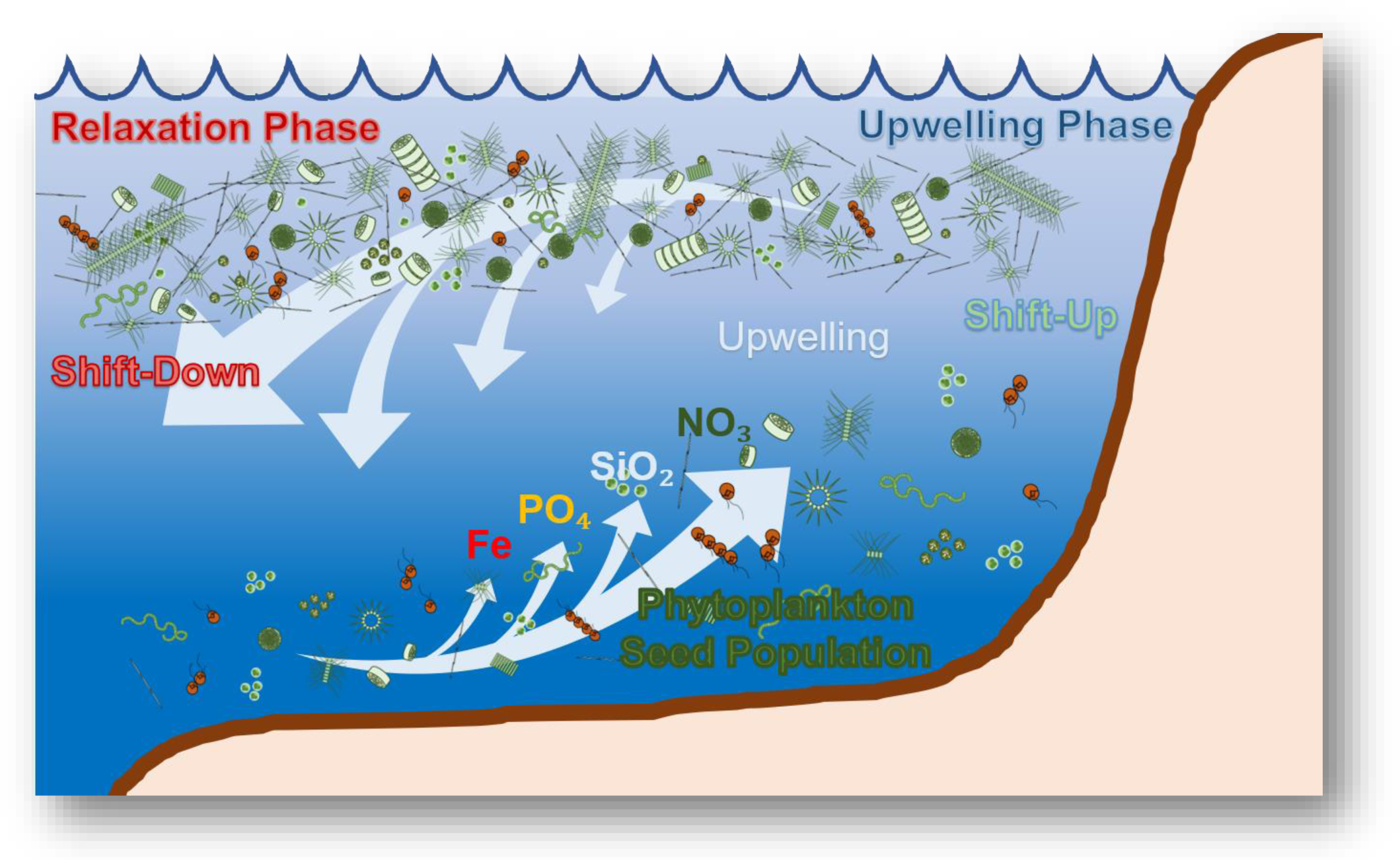
A conceptual model of the upwelling conveyor belt cycle (UCBC) in upwelling systems and the idealized zones within the upwelling cycle. Image modified from Lampe et al. (2021).

Upwelling-simulation studies conducted in the laboratory have recently showed an increase in the relative expression of nitrogen assimilation genes in the dark (simulated sinking out of the euphotic zone) in the diatom *Chaetoceros decipiens*, while the coccolithophore, *Emiliania huxleyi* showed a relatively lower expression of the same genes in the dark, further supporting the diatom “frontloading” hypothesis for N assimilation [Lampe et al. 2021]. Ultimately, diatoms can ensure a more rapid physiological response to upwelling compared to other phytoplankton taxa. Once they are upwelled, diatoms may rapidly increase growth through enhanced nitrogen metabolism and continue building up photosynthetic machinery until nitrate depletion or sinking. This highly dynamic series of physical and biological phases is defined as the upwelling conveyor belt cycle.

In essence, upwelled diatoms employ a proactive approach and maintain elevated pools of these nitrogen-related genes throughout the upwelling conveyor belt cycle, whereas other phytoplankton employ a more reactive approach and upregulate primary nitrate assimilation genes post-upwelling. This distinctive molecular strategy to constitutively express nitrogen assimilation genes is believed to provide these diatom taxa with a physiological edge in their “shift-up” response to upwelling [Lampe et al. 2021].

Iron (Fe) is yet another nutrient that differentially influences phytoplankton growth and community composition. It is an important micronutrient found in the photosynthetic reaction centers (photosystems I and II), and is vital to both oxygenic photosynthesis and ATP synthesis [Borowitzka et al. 2016, Falkowski 1997]. Furthermore, Fe plays a significant role in nitrogen assimilation, where the assimilatory enzymes nitrate reductase (NR) and nitrite reductase (NiR) both contain Fe. Fe is a limiting nutrient to productivity in vast offshore regions of the ocean, as phytoplankton subjected to low Fe availability have both experienced reductions in photosynthetic rates as well as nitrite reductase activity [Milligan and Harrison 2000]. Fe bioavailability further influences Fe uptake mechanisms in phytoplankton. Most of the dissolved Fe is complexed to organic ligands [Gledhill et al. 2012, Hutchins and Boyd 2016], but a small amount of the unchelated labile Fe(III) - or Fe’ - is found to be the most readily available source of Fe for phytoplankton [Turnšek et al. 2019].

In response to Fe-limited growth conditions, marine diatoms may produce Fe-starvation-induced-proteins (ISIP1, ISIP2A/pTF, ISIP3) [Cohen et al. 2018b, Gao et al. 2021. Turnšek et al. 2019] to facilitate Fe acquisition, including the endocytosis of Fe-bound siderophores [Kazamia et al. 2018]. ISIP2A has recently been identified to function as a phytotransferrin (pTF) in diatoms [McQuaid et al. 2018]. The significance of these phytotransferrins to algae isn’t exclusively limited to Fe trafficking, however, as previous studies have established a synergistic interaction of labile Fe and carbonate ions within the Fe uptake system, where phytotransferrin internalization via endocystosis is mediated by a tight coupling with the carbonate anion [McQuaid et al. 2018]. Effectively, the carbonate anion is protonated while the ferric Fe bound to the transferrin is reduced, which coordinates the release of Fe into the cell post-endocytosis [Baker et al. 2003]. Given that future acidification will likely decrease the carbonate ion concentrations in seawater, this could pose severe consequences on high-affinity Fe acquisition in diatoms. Yet, as diatom resilience has been a key factor in their success for nitrogen assimilation [Lampe et al. 2018], their susceptibility to Fe limitation can be similarly parried by their notable opportunism for Fe acquisition and storage. Diatoms are known to outperform other phytoplankton taxa during Fe enrichment experiments [Coale et al. 1996]. Pennate diatoms such as *Pseudo-nitzschia* have been shown to utilize the Fe-concentrating protein ferritin (FTN) during Fe-replete conditions to store excess Fe and maintain growth during periods of chronic Fe deficiency [Marchetti et al. 2009]. In other cases, the diatom proton-pumping protein rhodopsin (RHO) has also been implicated to help diatoms cope with Fe stress, where its presence in diatoms has allowed for a potential low-Fe requiring energy production alternative to photosynthesis [Cohen et al. 2017, Marchetti et al. 2015]. Nevertheless, the potential for Fe-limitation in future oceans should not be underestimated, where certain diatoms have been found to exhibit enhanced Fe-acquisition and storage strategies, others that thrive at present day Fe concentrations may not have the ability to compete in environments where a decline in bioavailable Fe occurs directly or indirectly due to climate change.

This study aims to understand the complexity of the phytoplankton response to the upwelling conveyor belt cycle, and simultaneously examine their physiological and transcriptomic changes in relation to Fe status within the CCS. The primary focus of this research is predicated on how Fe limitation can change and shape natural phytoplankton community structure, and how it plays a role in their acclimation to the upwelling conveyor belt cycle with the understanding that future coastal oceans are likely to experience dramatic shifts in Fe bioavailability across different topographical regions. Three central questions are addressed: (1) How does Fe limitation change the phytoplankton community structure, physiological response to upwelling, and gene expression of different phytoplankton groups during an upwelling event? (2) In what ways are diatoms responding to fluctuations in Fe bioavailability with respect to the upwelling conveyor belt cycle and topographical variation? (3) How much impact do diatom seed populations impose on their ability to respond to an upwelling event, and what are the biological driving forces that aid in their acclimation to upwelling?

## Methods

### Experimental Design

The primary objective of this study was to observe how subsurface phytoplankton communities respond to the upwelling conveyor belt cycle. To capture this phenomenon, we conducted on-deck incubations that spanned different time points which represented the different stages of growth (initial deep-water community at T0, and stimulated growth phases at T1 and T2). We further subjected the incubations to various Fe-related treatments to test how Fe might play a role in determining their physiological and molecular response to upwelling. Ultimately, the incubations allowed us to examine and track the growth of the CCS phytoplankton communities through a simulated upwelling experiment.

### Sample Sites

The first incubation was conducted from May 27^th^ to June 1^st^ of 2019. The site (41°0’52.956” N, 124°24’58.68” W) was located off the coast of northern California over a wide continental shelf (Supplemental Fig. 1). Upwelling conditions were not present at this site during the initial collection for the incubation experiment on May 27^th^. Sea surface temperature (SST) data obtained from remote sensing before, during, and after this wide shelf incubation suggests that upwelling had ceased approximately 11 days prior to the collection, and the site was in a period of relaxation throughout the incubation experiment (figure 2).

**Figure 2.**
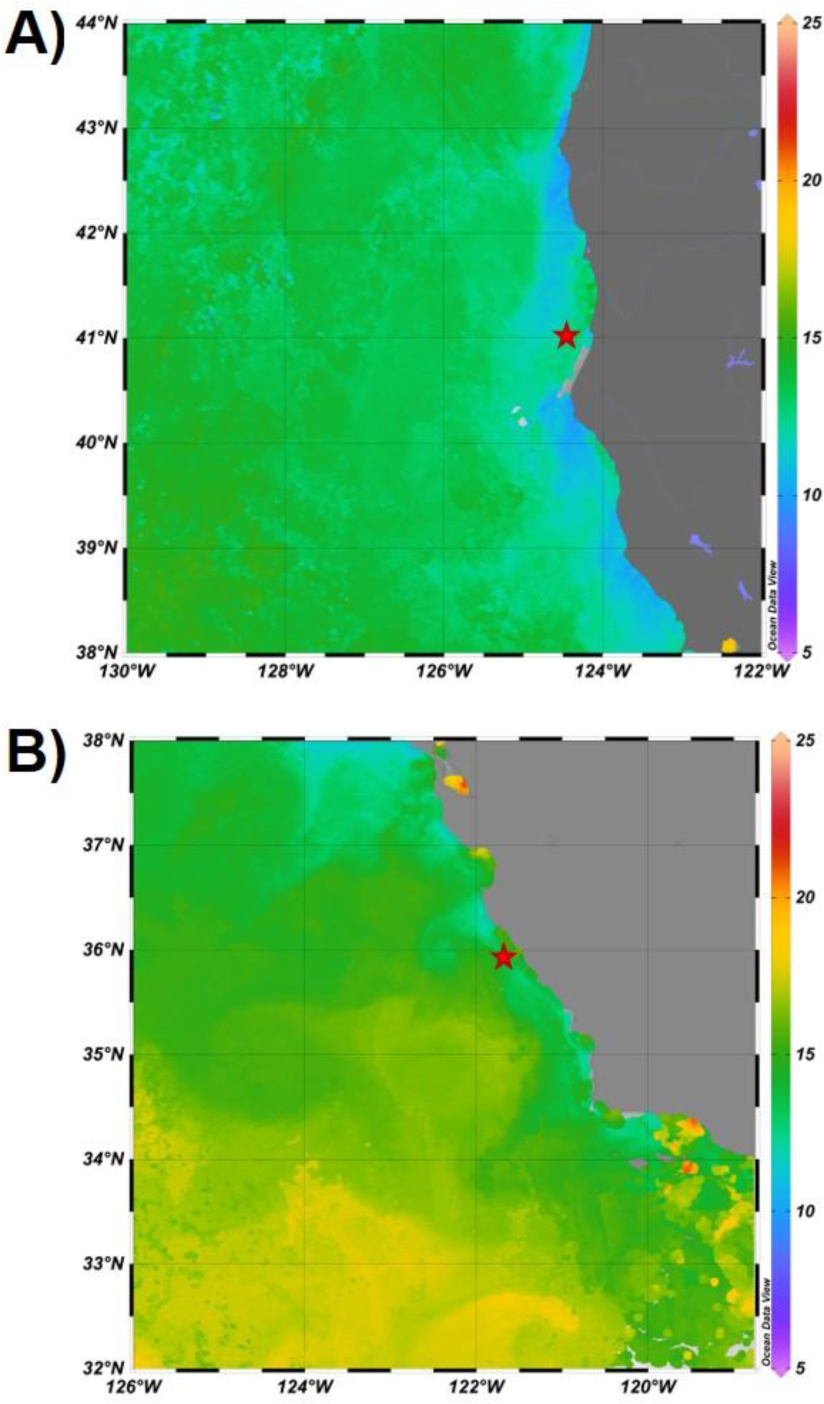
Remote sensing imagery (14-day averages) of sea surface temperature in the A) wide shelf and B) narrow shelf regions at the time of collection. Red stars on each map denote the respective location of deep-water collection for the incubation experiments. Remote-sensing data was derived from NOAA CoastWatch program.

The second incubation was conducted from June 2^nd^ to June 6^th^ of 2019. The site (35°55’24.7074” N, 121°32’34.44” W) was located in the Big Sur region of central California and characterized by a narrow continental shelf (Supplemental Fig. 1) where Fe is historically thought to be low [Bruland et al. 2001]. SST data for the narrow shelf incubation at Big Sur indicated that the region had not experienced an upwelling event for over a month prior to the experiment, but was transitioning to upwelling conditions prior to our occupation and was in an upwelling state during the incubation experiment (Fig. 2).

For this experiment, we simulated upwelling by pumping water from the isotherm range that would bring nutrient-rich cold water to the surface once upwelling favorable conditions were reinstated. Water temperatures within the 8-10 °C isotherms were used as the threshold. Satellite-derived data for sea surface temperature was obtained from NOAA POES AVHRR satellite (NOAA/NESDIS Center for Satellite Applications Research), downloaded from the NOAA CoastWatch Browser and plotted with GraphPad Prism v9.2.0.

### Water collection and incubation

Seawater was collected using trace-metal clean techniques from a depth of 90 m (corresponding to the 8.4 °C isotherm) for the wide shelf incubation and 80 m (corresponding to the 8.9 °C isotherm) for the narrow shelf incubation. Seawater from both sites was pumped directly into a positive pressure trace metal clean plastic bubble created in the ship’s laboratory into large, 50-gallon acid-washed high-density polyethylene (HDPE) drums to homogenize the seawater using a Wilden air-operated double-diaphragm pump made of polytetrafluoroethylene (PTFE) and acid-washed HDPE tubing. Preparation of the cubitainers was carried out prior to the cruise and included trace-metal clean techniques [Crawford et al. 2003]: cubitainers were initially soaked in 2% Extran detergent for 7 days, then rinsed with deionized water four times and Milli-Q water three times prior to being soaked in 10% reagent grade hydrochloric acid for 2-3 days and rinsed with Milli-Q water. Subsequently, the cubitainers were soaked in 1% trace metal grade hydrochloric acid for 7 days and rinsed with Milli-Q water, then soaked in 0.1% trace metal grade acetic acid 3-4 days before final storage in low Fe water from Station P (50 °N, 145 °W). Station P water had been collected and filtered from the 2018 EXPORTS North Pacific field campaign using trace metal clean techniques and stored in the dark at 4 °C until use.

In the bubble, these triplicate acid-cleaned 10 L low-density polyethylene (LDPE) cubitainers were filled and then incubated in large on-deck plexiglass incubators circulated with water chilled to the temperatures at which the subsurface samples were collected using Aqua Logic Delta Star® In-Line Water Chillers. Incubators were covered with neutral density screening to achieve 30% incident irradiance. To assess the effects of Fe addition or removal on the simulated upwelled plankton communities, samples were incubated with no amendment (control), amended with 5 nM of FeCl_2_ (Fe treatment), or amended with 200 nM Desferrioxamine B (DFB), a strong Fe chelator which inhibits dissolved Fe uptake (DFB treatment). Three cubitainers were immediately harvested for the initial timepoint (T_0_), also referred to as deep-water or DW. The remaining cubitainers were incubated for two additional timepoints for a total of 18 cubitainers per incubation. Cubitainers were harvested for various biological and chemical parameters (see below) following 48 hours (T_1_) for both incubation experiments, and following 120 hours in the wide shelf incubation and 96 hours in the narrow shelf incubation (T_2_).

### Physiological measurements

For chlorophyll *a* measurements, 250 mL of seawater was gravity filtered through 5 μm Isopore membrane filters (47 mm) and subsequently vacuum filtered onto GF/F (25 mm) filters under 100 mmHg of vacuum pressure. Filters were then rinsed with 0.45 μm filtered seawater and immediately stored at -20 °C until onshore analysis in the lab. Chlorophyll *a* extraction was performed using a 90% acetone solution at -20 °C for 24 hours and measured on a 10-AU fluorometer (Turner Designs, San Jose, CA) using the acidification method [Parsons et al. 1984].

Cell abundances were determined via flow cytometry. Two mL of whole seawater sample was collected and preserved in 10% paraformaldehyde solution (PFA, 1% formaldehyde) at -80 °C. Prior to flow cytometry analysis, the samples were thawed at room temperature. One mL of each whole seawater sample was transferred to a clean 5 mL Falcon Round-Bottom Polystyrene test tube (ThermoFisher Scientific™). The volume of the sample was measured as the difference between the mass of the tube before and after flow cytometry processing divided by the density of seawater. The samples were then processed through the BD FACSMelody™ (NC State University), which was set to an excitation wavelength of 488 nm and an emission peak of 670 nm. Phytoplankton functional groups were determined and gated (Supplemental Fig. 2) by their relative emission spectra of chlorophyll *a* (PerCP) as a dependent function of their size, plotted as the forward scatter (FSC-H). The cyanobacterium *Synechococcus* was further distinguished by its phycoerythrin (PE-H) emission spectra as a dependent function of its side scatter (SSC-H). A microscopy check was conducted after an initial fluorescence-assisted cell sorting (FACS) of a test sample to confirm the precision of the gated populations, which led to the determination of the following functional groups: large centric diatoms, small diatoms, picoeukaryotes, *Synechococcus*, and other (including dinoflagellates or diatom cells with low chlorophyll *a* content).

To assess the isotope uptake of DIC and NO_3_, a 618 mL polycarbonate bottle was filled to the top with incubation samples from each cubitainer. The subsamples were collected for dissolved inorganic carbon (DIC) and nitrate uptake rates at each respective time point, and immediately spiked with both NaH^13^CO_3_ and Na^15^NO_3_ at approximately 10% of the estimated ambient DIC concentrations and measured nitrate concentrations in a trace metal clean (TMC) space located on the ship (see Supporting Information S.1.1 for more details). Since there were no underway measurements of DIC, an expected ambient concentration of 2000 μmol L^-1^ was derived from literature values for both incubation sites [Fassbender et al. 2011]. Nitrate concentrations of the deep-water and surface water in both incubation sites were measured using the Submersible Ultraviolet Nitrate Analyzer (SUNA), which quantifies dissolved nitrate concentrations by illuminating the water sample with UV and using the absorbance values to estimate nitrate concentrations from a multi-variable linear regression based on the MBARI method [Johnson and Coletti 2002]. After spiking with respective trace concentrations of both stable isotopes of DIC and nitrate, samples were returned to the incubator for six hours. All rate measurement incubations were initiated at approximately the same time of day (i.e., close to dawn). Following incubation, seawater was immediately filtered where cells greater than 5 μm were gravity filtered onto 5 μm Isopore membrane filters (47 mm) and then washed onto a pre-combusted (450 °C for 5 hours) GF/F filters (0.7 μm nominal porosity) by vacuum filtration using 0.2 μm filtered seawater and preserved at –20 °C. Cells smaller than 5 μm passed through the initial 5 μm Isopore membrane filters and were collected on pre-combusted GF/F filters and preserved at –20 °C. Prior to analysis, filters were dried at 60 °C for 24 hours, encapsulated in tin and pelletized. Particulate organic carbon (POC), particulate organic nitrogen (PON), and atom percentages of ^13^C and ^15^N were subsequently quantified using an isotope ratio mass spectrometer (EA-IRMS) at the UC Davis Stable Isotope Facility. For each sample, POC and PON concentrations (μmol L^-1^) were calculated by dividing the measured POC/PON mass (μg) by the respective atomic mass of carbon and nitrogen over the volume filtered (0.62 L). Absolute uptake rates (see Supporting Information S1.1) were derived from the constant transport model (Eq. 3 from Dugdale and Wilkerson 1986). Biomass-normalized uptake rates (*V*, DIC or NO_3_ taken up per unit POC or PON, respectively, per unit time) were derived from the specific uptake model (Eq. 6 from Dugdale and Wilkerson 1986), and was assessed by dividing the absolute uptake rates of DIC or NO_3_ by the measured POC and PON concentrations (μmol L^-1^), respectively. Correction for ammonium regeneration during the incubation period was not performed.

Dissolved inorganic nutrients (nitrate + nitrite, phosphate and silicic acid) were measured by filtering 30 ml of water through a 0.2 µm filter, using acid-washed syringes into an acid-cleaned polypropylene Falcon^TM^ tube. Dissolved nutrient concentrations were analyzed using an OI Analytical Flow Solutions IV auto analyzer by Wetland Biogeochemistry Analytical Services at Louisiana State University. Concentrations of nitrate measured from the discrete samples were used for the calculation of absolute nitrate uptake rates.

### RNA collection and meta-transcriptomic analysis

Approximately 2.5 L of seawater from each cubitainer was collected onto 0.8 μm Pall Supor filters (142 mm) using a peristaltic pump, then immediately flash frozen in liquid nitrogen, and stored at -80 °C until RNA extraction in the laboratory. RNA was extracted using the RNAqueous-4PCR kit. An additional bead beating step and higher volumes (3 mL) of lysis buffer and ethanol were added to the extraction procedure, which was followed according to the manufacturer’s instructions. The glass beads were used to assist in disrupting cells, particularly those with hard outer coverings (e.g., diatoms). RNA quantity and purity was assessed on a NanoDrop 2000 spectrophotometer (ThermoFisher Scientific™). Total RNA for the control, Fe, and DFB treatments at the 48hr time point (T_1_), and the DFB treatment at 120h time point (T_2_) in the wide shelf incubation were pooled into singular samples due to low RNA yields. All other biological replicates were maintained separately for sequencing. Library preparation and sequencing of RNA with poly-A tail selection was conducted by GENEWIZ. Sequencing was performed on an Illumina HiSeq 4000 with a 2×150 bp configuration.

Reads were trimmed for quality control and adapter removal using Trimmomatic v0.32 (paired-end mode, sliding window of 4:20, and minimum length of 36 bp) and FastQC(Babraham Institute) was used for quality control assessment. Trimmed pared reads were then assembled *de novo* into contigs using rnaSPAdes v3.14.1. The assemblies were then clustered using CD-HIT-EST to reduce redundant contigs [Li et al. 2006] based on a 98% similarity score, and merged into a mega assembly. Read counts were quantified by mapping trimmed reads to the assembled contigs using SALMON alignment [Patro et al. 2017].

Taxonomic and functional annotations were both performed using the DIAMOND sequencing aligner and its compatible BLASTX command (E-value < 10^-6^). Taxonomic annotation was identified through phyloDB (v1.076), a custom reference database that also includes data collected from the Marine Microbial Eukaryote Transcriptome Sequencing Project (MMETSP) [Keeling et al. 2014]. Functional annotation was assigned to contigs using the Kyoto Encyclopedia of Genes and Genomes (KEGG) [Kanehisa et al. 2017]. The best hit with a KEGG Ortholog (KO) number from the top hits was selected, and the KEGGANNOT package was used to obtain annotation data for contigs with assigned KOs. Contigs of importance such as those for the Fe starvation induced proteins (*ISIP*s) and proton-pumping rhodopsin (*RHO*) without assigned KOs were manually added into the subsequent KEGG BLASTX output with a “pseudo” KO identifier.

Prior to differential expression analysis, read counts were imported and summarized into gene-level estimates for each respective taxonomic group (diatom, dinoflagellate, chlorophyte, haptophyte) and diatom genus (*Chaetoceros*, *Pseudo-nitzschia*, *Thalassiosira*) using Tximport (v3.13) [Soneson et al. 2016]. Differential expression analysis was performed using DESeq2 [Anders and Huber 2010], and assessed by summing counts of contigs within different taxonomic groups and diatom genera, respectively. The read counts were normalized within each group and genus and by the respective incubations (wide shelf and narrow shelf) using the “median ratio method” (Eq. 5, Anders and Huber 2010), and dispersions were estimated from treatments where replicate samples were individually sequenced using a negative binomial distribution. For hypothesis testing between two groups, the Wald test was used by dividing log_2_ fold change by its standard error and comparing the resulting z-statistic to a standard normal distribution. A p-value of less than 0.05 would thus indicate significant differential expression. Genes displayed for volcano plots and heatmaps were filtered by a log_2_ fold change cutoff of greater than 1 or less than -1 and a p-adjusted value of less than 0.05. Plots were created using the program R v4.0.1 on RStudio.

### Statistical Analyses

Two-way ANOVAs followed by Tukey tests for multiple comparisons were performed on the physiological uptake rate properties of the incubation treatments (non-gene expression data) using the SciPy and Statsmodels packages in Python 3.11.2. Three comparisons were made between the different treatment groups (control and Fe, control and DFB, Fe and DFB). For response variables, absolute and biomass-normalized uptake for DIC and nitrate within the large (≥5 μm) or small (<5 μm) cells in either the wide or narrow shelf were summarized with the time points of the incubations (T_0_, T_1_, and T_2_), and compared relative to the predictor variables (comparisons between treatments). The Bioinfokit toolkit was used to conduct the Tukey tests and produce a Tukey summary. The main output of the ANOVAs is discussed in Supporting Information S2.2.

### Data deposition

The sequence data reported in this study will be deposited to the National Center for Biotechnology (NCBI) sequence read archive. RNA sequences are under submission no. SUB13179044 (Bioproject accession no. PRJNA966115), and rDNA sequences are under the submission no. SUB12523058 (Bioproject accession no. PRJNA966115). Assembled contigs, read counts, and annotations are deposited to Zenodo. Data for this project was also submitted to BCO-DMO under project number 768006.

## Results

### Shelf characteristics during water collections

The ambient deep-water communities (DW) incubated for the simulated upwelling experiments were collected at roughly 8.4 °C isotherm for the wide shelf (90 meters) and 8.9 °C isotherm for the narrow shelf (80 meters). For the wide shelf seawater, sea surface temperature (SST) data derived from satellite imagery provided evidence that upwelling had ceased 11 days prior to the start of the on-deck incubation, and the site was in a state of relaxation throughout the remainder of the experiment. In contrast, SST data for the narrow shelf suggested that the area was transitioning to upwelling conditions just prior to our occupation, and was in an upwelling state during the on-deck incubation (Fig. 2D).

### Macronutrient concentrations and drawdowns

Macronutrient concentrations in both incubations remained high throughout the incubation, and there is no evidence of macronutrient limitation or depletion at any point during the incubation experiments. Initial NO□ (16.1 ± 1.65 µmol L^-1^), PO□ (1.73 ± 0.14 µmol L^-1^), and Si(OH_4_) (22.6 ± 3.77 µmol L^-1^) concentrations in the wide shelf were higher than the initial NO□ (12.7 ± 1.62 µmol L^-1^), PO□ (1.50 ± 0.14 µmol L^-1^), and Si(OH)_4_ (15.8 ± 2.47 µmol L^-1^) concentrations found in the narrow shelf. Furthermore, no appreciable decline in macronutrient concentrations were detected in any of the treatments by T_2_ in the wide shelf incubations (Fig. 3). Differences in macronutrient concentrations in the wide shelf incubation across the different time points were found to be insignificant (p > 0.05) amongst all three treatments (Supplemental Table 1). It is also likely that any apparent increases in macronutrient concentrations in the wide shelf incubation could be the result of cellular remineralization from the lower growth rates in the DFB treatments relative to the control and Fe treatments, suggesting that there was slight heterogeneity amongst the different cubitainers. A drawdown of NOL, POL, and Si(OH)_4_ at T_2_ in the narrow shelf incubations was observed, corresponding to an increase in biomass at the same timepoint for both the control and the Fe treatments (see below).

**Figure 3.**
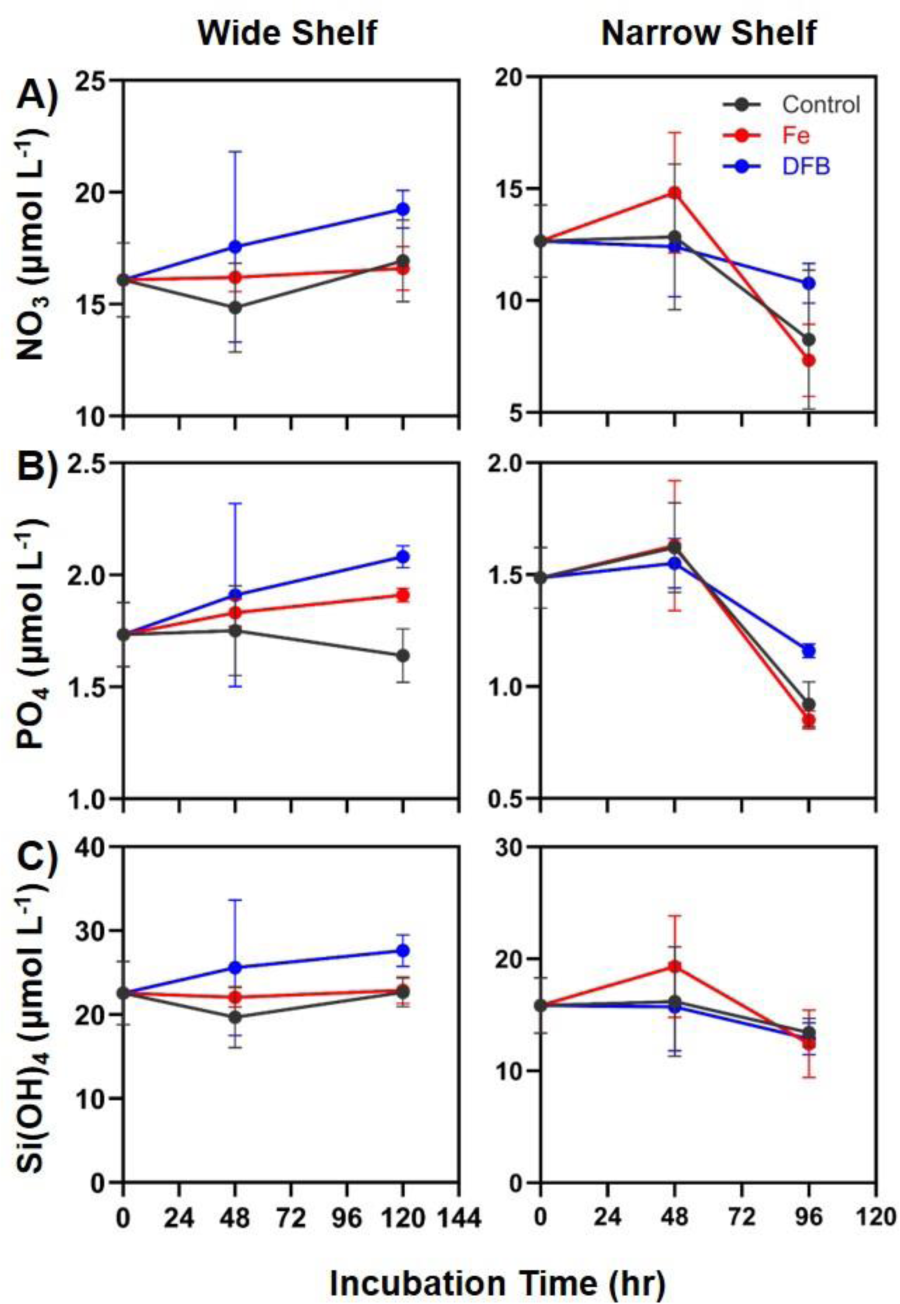
Macronutrient concentrations of A) nitrate (NO3), B) phosphate (PO4), and C) silicic acid (Si[OH]4) of the wide shelf and narrow shelf incubations. Black, red and blue symbols and lines represent the control, Fe, and DFB treatments, respectively. Error bars represent the standard deviations of the mean (n=3).

### Phytoplankton biomass

Chlorophyll *a* concentrations were generally lower in the wide shelf relative to that of the narrow shelf incubations. Comparing across sites, the wide shelf incubation started with a relatively lower chlorophyll *a* biomass (0.019 ± 0.003 µg L^-1^, average ± standard deviation) at T_0_ compared to that of the narrow shelf incubation (0.31 ± 0.05 µg L^-1^) for the large size-fraction (Supplemental Table 2). Similarly, the initial deep-water community of the small size-fraction in the wide shelf incubation (0.015 ± 0.003 µg L^-1^) was five-fold lower in biomass than that of the narrow shelf (0.10 ± 0.01 µg L^-1^). In the wide shelf incubation, chlorophyll *a* data indicated an increase of large phytoplankton (≥5 µm) by T_2_ in the control (0.66 ± 0.14 µg L^-1^) and Fe (0.80 ± 0.26 µg L^-1^) treatment, as the DFB treatment (0.08 ± 0.04 µg L^-1^) yielded observably lower biomass relative to the control and Fe treatments. For the smaller phytoplankton (<5 µm), the control (0.12 ± 0.09 µg L^-1^), Fe (0.09 ± 0.04 µg L^-1^), and DFB treatment (0.04 ± 0.02 µg L^-1^) were markedly lower in biomass by T_2_ than those of the larger fraction. In the narrow shelf incubation, chlorophyll *a* data also indicated an increase of mostly large phytoplankton by T_2_ of the incubation in the control (10.7 ± 4.1 µg L^-1^, average ± standard deviation), Fe (8.2 ± 1.4 µg L^-1^), and DFB (3.31 ± 0.47 µg L^-1^) treatments, although DFB had a noticeable negative effect on biomass accumulation. Differences in chlorophyll *a* for the smaller phytoplankton among the control (0.59 ± 0.28 µg L^-1^), Fe (1.16 ± 0.35 µg L^-1^), and DFB (0.79 ± 0.07 µg L^-1^) treatments at T_2_ of the narrow shelf incubation are less distinct. Interestingly, DFB did not seem to appreciably influence biomass accumulation of the smaller phytoplankton (Fig. 4B).

**Figure 4.**
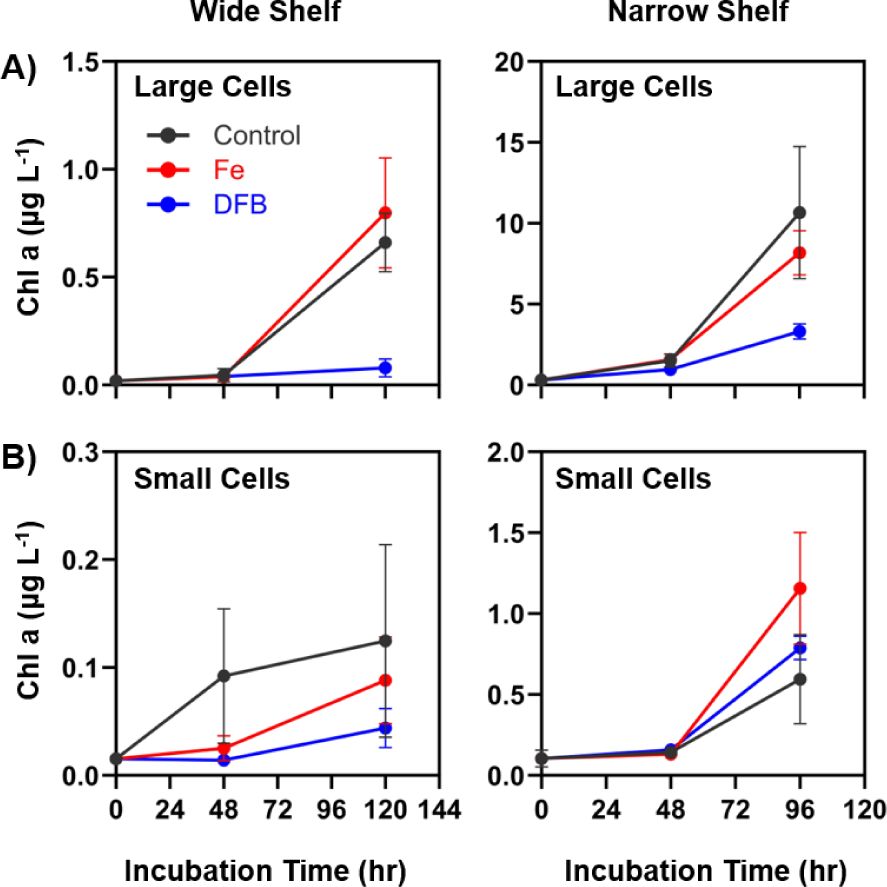
Chlorophyll *a* concentrations over the time course of the wide shelf and narrow shelf incubations for the A) large cell size-fractions (≥5 μm) and for the B) small cell size-fraction (<5 μm). Black, red and blue circles and lines represent values of the control, Fe and DFB treatments, respectively. Chlorophyll *a* concentrations for the initial deep-water seed communities (T0) are provided in Supplementary Table 1A. Error bars represent the standard deviations of the mean (n=3).

In the wide shelf incubations, cell counts (Fig. 5A) derived from fluorescence-assisted flow cytometry at T_2_ show a pronounced increase in diatoms (small diatoms and large centric diatoms) in the control (994 ± 33.6 cells mL^-1^, small diatom; 184 ± 7.1 cells mL^-1^, large centric) and Fe treatments (1005 ± 213.4 cells mL^-1^, small diatom; 209 ± 78.9 cells mL^-1^, large centric), as DFB had a noticeable inhibiting effect on diatom growth (270 ± 15.2 cells mL^-1^, small diatom; 31.8 ± 4.2 cells mL^-1^, large centric). Neither the small diatoms or the large centric diatoms displayed any noticeable differences in abundance across the treatments at T_1_ of the wide shelf incubation (Fig. 5A). The pico-eukaryotes displayed similar patterns to that of the diatoms in the control (66.4 ± 5.2 cells mL^-1^, T_1_; 306.8 ± 110.7 cells mL^-1^, T_2_), Fe (45.7 ± 21.4 cells mL^-1^, T_1_; 213.4 ± 35.6 cells mL^-1^, T_2_), and DFB (8.9 ± 4.1 cells mL^-1^, T_1_; 18.9 ± 7.9 cells mL^-1^, T_2_) treatments, but were noticeably lower in abundances relative to the diatoms. The cyanobacteria *Synechococcus* were found to be more abundant at T_0_ in the wide shelf (92.2 ± 3.6 cells mL^-1^) relative to the rest of the incubation time points, but also displayed DFB effects at both T_1_ and T_2_ (Fig. 5A). In the narrow shelf, no noticeable difference in small diatom abundances across treatments was observed at T_1_ or T_2_, although DFB (636 ± 56.5 cells mL^-1^) had a negative effect on the population average of large diatoms relative to the control (2690 ± 20.9 cells mL^-1^) and Fe treatments (2977 ± 390.9 cells mL^-1^) at T_2_ (Fig. 5B). Similar to that of the large diatoms, the population average of pico-eukaryotes in the narrow shelf incubation also displayed a DFB effect (927 ± 71.0 cells mL^-1^) relative to the control (2827 ± 807.1 cells mL^-1^) and Fe treatments (3834 ± 189.5 cells mL^-1^) at T_2_, while no noticeable difference was observed at T_1_. The *Synechococcus* population was also found to be more abundant at T_0_ (250 ± 8.9 cells mL^-1^) compared to the rest of the incubation time points, but did not show a DFB effect at T_2_ (Fig. 5B).

**Figure 5.**
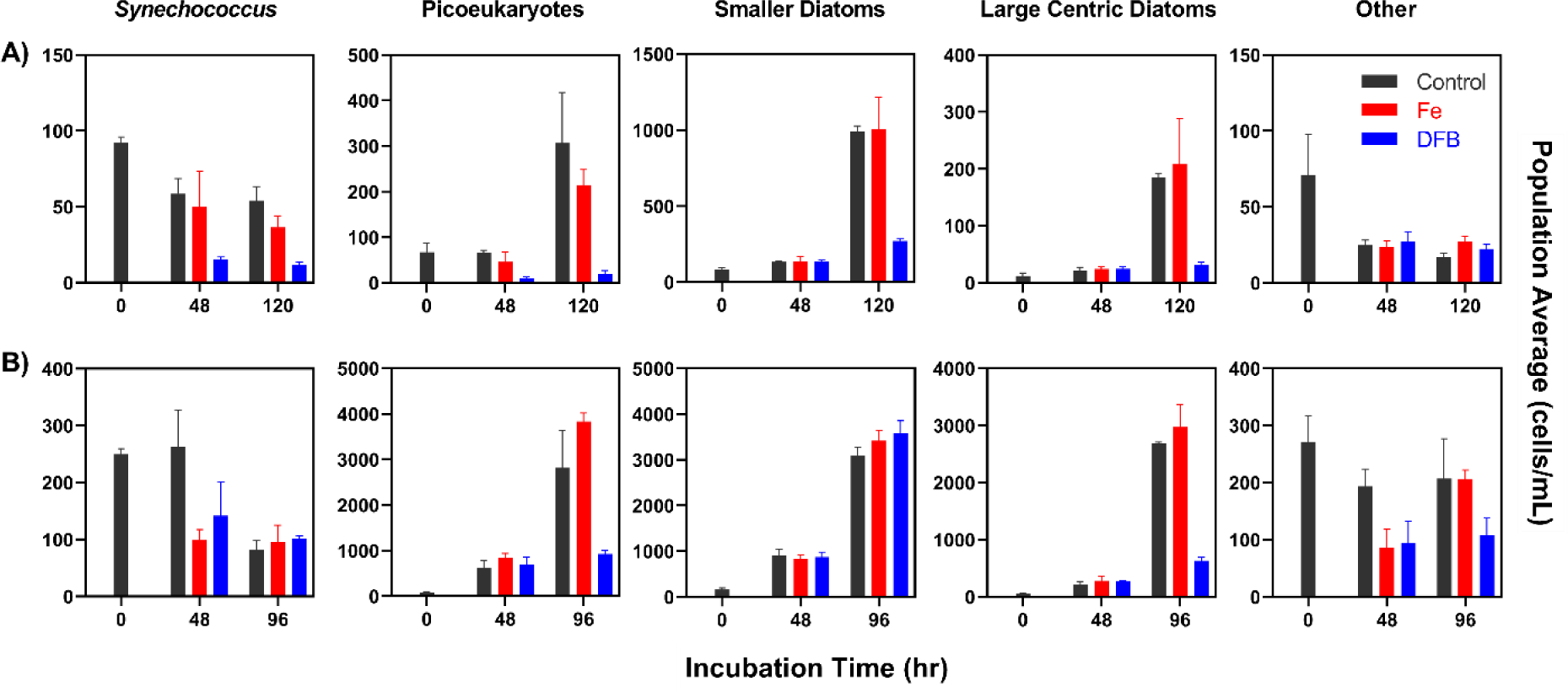
Flow cytometry (FCM) derived cell densities (cells mL^-1^) at each incubation time (hours) of different phytoplankton populations in response to the control, Fe, or DFB treatments in the A) wide shelf incubation experiment and B) narrow shelf incubation experiment. Averages and standard deviation were measured from triplicate treatments. Grey, red, and blue bars represent population averages from control, Fe, and DFB treatments, respectively. Population averages for the initial seed community at T0 are provided in Supplementary Table 1B. Growth rates derived from FCM counts of pico-eukaryotes, smaller diatoms, and large centric diatoms are provided in Supplementary Table 2. Error bars represent the standard deviations of the mean (n=3).

### Inorganic carbon and nitrate uptake rates

Biomass-normalized DIC (0.0018 ± 0.001 hr^-1^) and nitrate (0.0038 ± 0.0019 hr^-1^) uptake of the large cell (≥5 µm) deep-water (T_0_) community in the wide shelf incubations (Fig. 6A) were lower than the DIC (0.0105 ± 0.0023 hr^-1^) and nitrate uptake rates (0.008 ± 0.002 hr^-1^) in the narrow shelf incubations. For deep-water (T_0_) phytoplankton <5 µm, biomass-normalized DIC uptake (Fig. 6B) in the wide shelf incubation (0.0013 ± 0.0007 hr^-1^) was lower than that of the narrow shelf incubation (0.0044 ± 0.0013 hr^-^1). Biomass normalized nitrate uptake rates for both the wide (0.0041 ± 0.002 hr^-1^) and narrow (0.0036 ± 0.0011 hr^-1^) shelf incubations were comparable in the small cell (<5 µm) T_0_ community. Absolute uptake DIC and nitrate uptake rates (Supplemental Fig. 3) are discussed in the supporting information.

**Figure 6.**
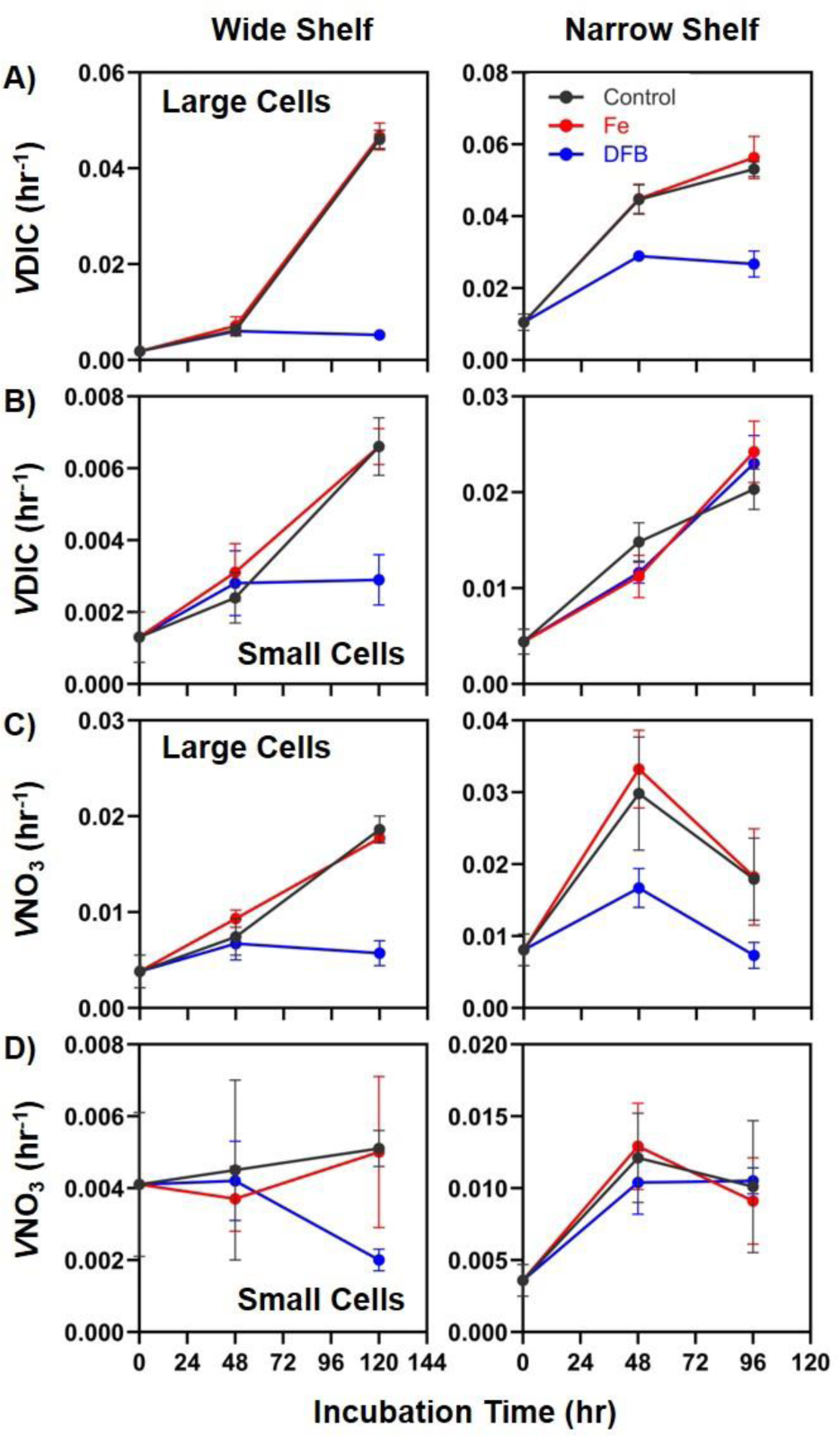
Biomass-normalized (*V*) specific uptake rates of dissolved inorganic carbon (DIC) in the A) large cell size-fractions (≥5 μm) and B) small cell size-fractions (<5 μm), and biomass-normalized (*V*) specific uptake rates of nitrate (NO₃) in the C) large cell size-fractions and D) small cell size-fractions over the course of the incubation at the wide and narrow shelves. Biomass-specific uptake rates are normalized to respective particulate carbon and nitrogen within the size fractions and treatments. Black, red and blue symbols and lines represent the control, Fe, and DFB treatments respectively. Error bars represent the standard deviations of the mean (n=3).

In the wide shelf incubation, biomass-normalized DIC and nitrate uptake rates of the larger phytoplankton (≥5 µm) did not exhibit significant differences among any treatment variable at T_1_, but show accelerated growth by T_2_ in the control and Fe treatments, while the DFB treatment exhibited considerably lower DIC uptake by T_2_ (Fig. 6A). Additionally, nitrate uptake rates in the control and Fe treatment were significantly greater than that of the DFB treatment (Fig. 6C). In the narrow shelf incubation, biomass-normalized DIC uptake data suggest that DIC uptake rates were starting to plateau by T_2_ in the large size-fraction (Fig. 6A). Interestingly, biomass-normalized nitrate uptake rates indicate a decline in nitrate uptake rate in the large size-fraction by T_2_, in which the DIC uptake rates had also plateaued (Fig. 6C). For the small size-fraction, no noticeable Fe or DFB effect on normalized nitrate uptake could be observed in the control (Fig. 6D)

In both the wide and narrow shelf incubations, statistical analysis of DIC and nitrate uptake rates (both absolute and normalized rates) of large phytoplankton (≥5 µm), based on post-hoc ANOVA tukey tests, suggest significant differences between control and DFB treatments (p < 0.05) and between Fe and DFB treatments (p < 0.05). No significant differences in rates (absolute and normalized) were observed between the control and Fe treatments in either cell size-fraction (Supplemental Table 1). Of the small cell size-fraction, statistical significance was observed in the comparison between Fe and DFB treatments for normalized DIC uptake (p < 0.05), but no statistical significance was found in any other comparison for either DIC or nitrate uptake (Supplemental Table 1).

### Taxonomic Composition

Taxonomic distribution based on normalized RNA transcript counts indicate diatom blooms under simulated upwelling in both incubations. The initial community in the wide shelf was comprised of a large proportion of dinoflagellates (47%) and other eukaryotes (35%), while diatoms (13%), chlorophytes (2%), and haptophytes (3%) were relatively low in abundance. However, diatoms represent a majority of the community in all treatments at T_1_ (control: 47%; Fe: 77%; DFB: 51%) and T_2_ (control: 69%; Fe: 88%; DFB: 60%), and Fe and DFB had differential effects on their relative proportion through the incubation (Fig. 7A).

**Figure 7.**
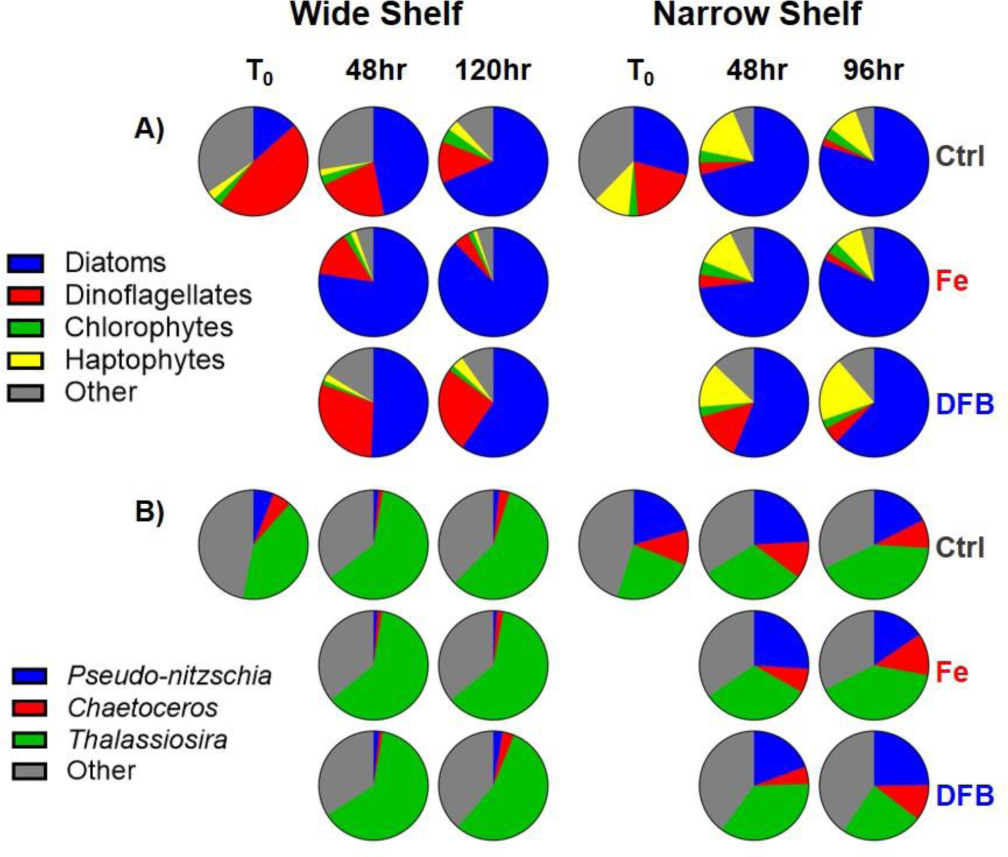
Average taxonomic distribution by normalized mapped RNA reads from each time point (T0: deep-water; T1: 48hr; T2: 120hr/96hr) and treatment (Control, Fe, and DFB, respectively). Taxonomic distributions are categorized as A) percentage of mapped reads from the whole phytoplankton community and B) percentage of mapped reads for the dominant diatom genera within all reads assigned and subset for diatoms (Bacillariophyta).

The initial community in the narrow shelf showed a stark contrast to that of the wide shelf in relative phytoplankton proportions, as there is a more even distribution of different phytoplankton taxa. Diatoms (29%) represent a higher proportion here compared to their counterparts in the wide shelf, while dinoflagellates (20%), haptophytes (11%), and chlorophytes (3%) comprise the rest of the main algal taxa. A large proportion of other eukaryotes (38%) could also be observed in the initial community (Fig. 7A). Diatoms similarly exhibit higher relative proportions in all treatments by T_1_ (control: 71%; Fe: 73%; DFB: 56%) and T_2_ (control: 80%; Fe: 82%; DFB: 62%), but the negative DFB effect on their transcript abundances are more noticeable compared to the control and Fe treatment at both time points (Fig. 7A).

Within the diatom taxa, *Thalassiosira* species (42%) represented the highest proportion of diatoms in the initial community in the wide shelf, while the genera *Pseudo-nitzschia* (6%) and *Chaetoceros* (5%) were the next most abundant. By T_1_ and T_2_, *Thalassiosira* maintained high relative proportions in all treatments (Fig. 7B). Similarly, in the narrow shelf initial community, *Thalassiosira* show the highest relative proportions of any other diatom genera (24%), but only marginally, as initial populations of *Pseudo-nitzschia* (21%) and *Chaetoceros* (10%) exhibited higher relative abundance in the narrow shelf compared to the wide shelf. This relatively even distribution of diatom genera could be observed throughout the incubation in all treatments (Fig. 7B), with a noticeable effect on *Thalassiosira* relative abundance in the DFB treatment (24%) compared to the control (42%) and Fe (40%) at T_2_. Contrastingly, at this same time point in the narrow shelf incubation, *Pseudo-nitzschia* seemed to respond positively to the DFB treatment (25%) and increased their relative abundance compared to the control (18%) and Fe treatments (15%). From our observations, while *Thalassiosira* (centric) diatoms responded positively to control and Fe treatments overall, *Pseudo-nitzschia* (pennate) diatoms seemed to thrive under Fe-limitation of the DFB treatments.

Taxonomic distribution based on 18S ribosomal DNA similarly indicate diatom blooms under upwelling conditions in both incubations by T_2_, but the diatom seed community in the wide shelf comprised a much smaller proportion than their narrow shelf counterparts (Supplemental Fig. 5A). Thus, diatoms in the narrow shelf incubation were able to represent a large majority of the phytoplankton community by T_1_, while diatoms in the wide shelf incubation were only able to comprise a large majority at T_2_.

### Phytoplankton community gene expression

Normalized transcript abundances for the four major phytoplankton taxa (diatoms, dinoflagellates, chlorophytes, and haptophytes) indicate differential expression of Fe starvation, nitrogen assimilation, and photosynthesis genes between the Fe and DFB treatments and across T_0_ (DW), T_1_, and T_2_ time points in both the wide shelf and the narrow shelf (Fig. 8). Since transcript abundances were normalized within each respective taxon, comparisons were made across the timepoints of an incubation for each group, and not across the taxonomic groups. The Fe starvation induced protein 3 (*ISIP3*) was conserved through all four major taxa, and showed higher transcript abundances in DFB treatments throughout both incubations and across the taxa. However, the DFB effect is more noticeable in diatom taxa and in the narrow shelf communities relative to the wide shelf communities. Interestingly, all three of the nitrogen assimilation related genes: nitrate transporter (*NRT*), nitrate reductase (*NR*), and ferredoxin-nitrite reductase (*nirA*) were high in transcript abundances in diatoms throughout the incubation starting in the initial deep-water community (DW) for both the wide and narrow shelf, but is even higher in proportion in the narrow shelf DW than the rest of the incubation. Conversely, the same nitrogen-related genes show more nuance for the other three major taxa, and are at higher abundance at the later time points relative to their DW expression. In dinoflagellates, relative transcript abundance of *nirA* was found to be low throughout both incubations (Fig. 8). Among the photosynthesis genes, cytochrome (*petA*, *petC*), photosystem I subunits (*psaA*, *psaB*, *psaC*, *psaD*, *psaE*, and *psaF*), and photosystem II subunits (*psbA*, *psbC*, *psbD*, *psbE*) were relatively lower in transcript abundance compared to the nitrogen assimilation gene transcripts in all observed taxa with the exception of cytochrome b6-f complex Fe-sulfur subunit (*petC*). The narrow shelf transcript abundances of these photosynthesis genes also show relatively higher abundance compared to those of the wide shelf, and this is observed in all four of the phytoplankton taxa (Fig. 8).

**Figure 8.**
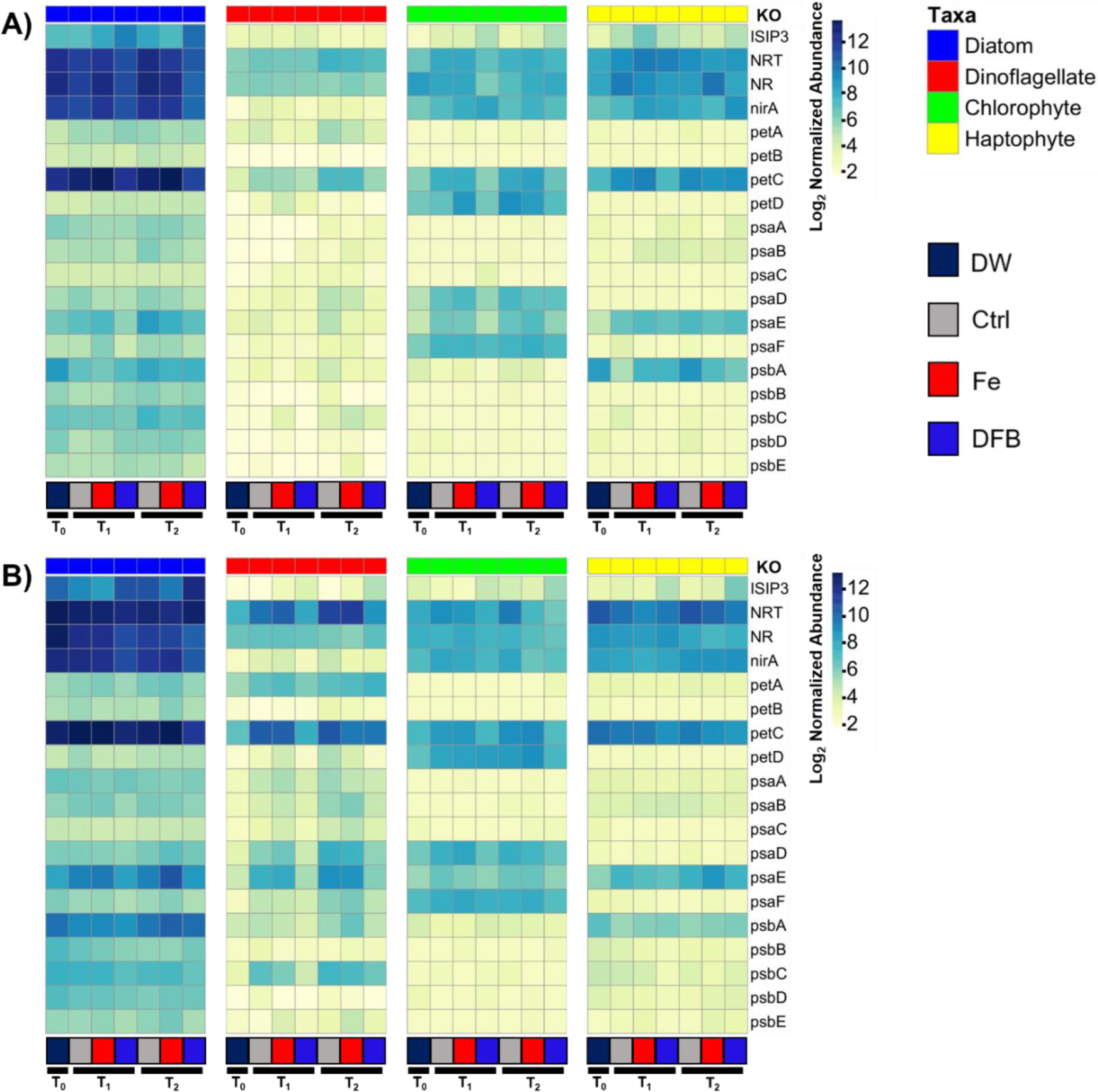
Normalized transcript abundances of select Fe starvation, Fe storage, nitrate assimilation, and photosynthetic genes identified amongst the four major phytoplankton taxa (diatoms, dinoflagellates, chlorophytes, and haptophytes) in the A) wide shelf incubation (T0 = DW, T1 = 48hr, T2 = 120hr) and B) narrow shelf incubation (T0 = DW, T1 = 48hr, T2 = 96hr). Abundances are normalized by total reads within each taxonomic group. The bottom of each heatmap represents the respective treatment and timepoint of the four different groups. Deepwater at T0 is denoted by DW, treatments at various time points are denoted by Ctrl, Fe and DFB.

### Diatom gene expression

The three major diatom genera (*Pseudo-nitzschia*, *Chaetoceros*, and *Thalassiosira*) exhibited higher relative abundances of *ISIP1*, *ISIP3*, and *pTF* genes in DFB treatments throughout both incubations, but ISIP1 expression was especially high in *Chaetoceros* and *Thalassiosira* at T_2_ in the DFB treatment in the narrow shelf (Fig. 9B). Ferritin (*FTN*) was proportionally abundant in the *Pseudo-nitzschia* and *Thalassiosira* genera throughout the incubation at both wide and narrow shelves, but a noticeably lower transcript abundance of *Thalassiosira FTN* can be observed in the DW of the narrow shelf compared to *FTN* transcript abundance in the DW of the wide shelf, while *Pseudo-nitzschia FTN* transcript abundance maintained its relative proportions in the DW of both incubation experiments (Fig. 9). Nitrogen assimilation gene transcripts in both incubations were higher in abundance in the DW, control, and Fe treatments relative to the DFB treatments for *Pseudo-nitzschia* and *Thalassiosira*; *NR* was observed to be highly expressed in the DW relative to the rest of the incubation in *Pseudo-nitzschia* and *Thalassiosira* genera of both incubation experiments. The DFB effect on nitrogen assimilation can be better distinguished at T_1_ of both incubation experiments for all three diatom genera. Similarly, the gene encoding cytochrome b6-f complex Fe-sulfur subunit (*petC*) exhibited higher fold change in the control and Fe treatments relative to the DFB treatments in all three diatom genera. Overall, higher transcript abundances for various photosynthesis genes can be observed in the narrow shelf incubation experiments, especially in *Thalassiosira* genera (Fig. 9B).

**Figure 9.**
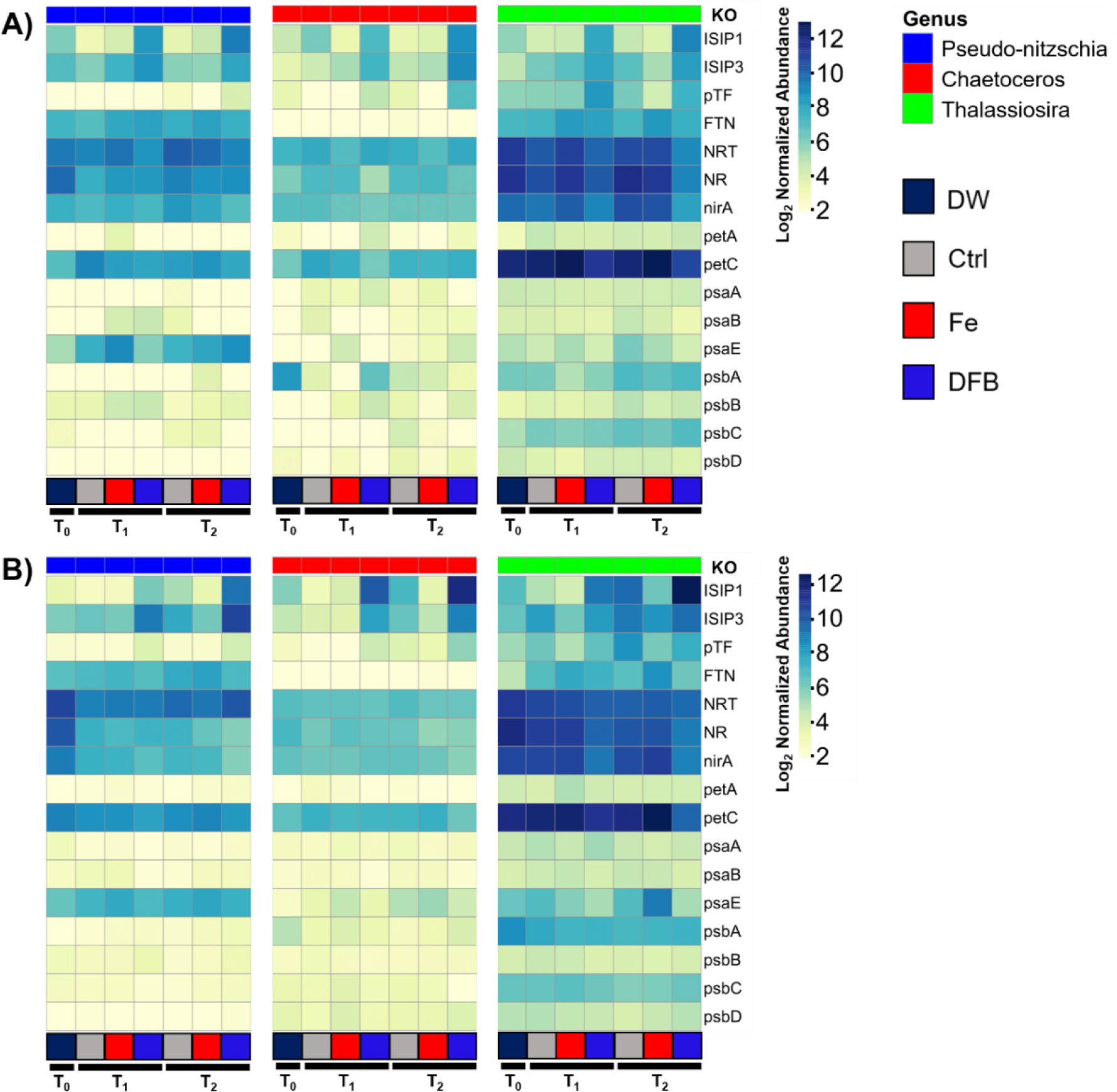
Normalized transcript abundances of select Fe starvation, Fe storage, nitrate assimilation, and photosynthetic genes identified amongst the three major diatom genera (*Pseudo-nitzschia*, *Chaetoceros*, and *Thalassiosira*) in the A) wide shelf incubation (T0 = DW, T1 = 48hr, T2 = 120hr) and B) narrow shelf incubation (T0 = DW, T1 = 48hr, T2 = 96hr). Abundances are normalized by total reads within each diatom genera. The bottom of each heatmap represents the respective treatment and timepoint of the three different genera. Deepwater at T0 is denoted by DW, treatments at various time points are denoted by Ctrl, Fe and DFB.

Diatom rhodopsin genes (*RHO*) exhibited significantly increased expression in the DFB treatments at both time points in the narrow shelf (p-adj < 0.05). Diatom *ISIP1*, *ISIP3*, and *pTF* genes were significantly differentially expressed between the Fe and DFB treatments at both time points in the narrow shelf incubation, and at T_2_ in the wide shelf incubation (p-adj < 0.05). *FTN* transcripts were also more abundant in the narrow shelf Fe treatments relative to the DFB treatments at both time points (p-adj < 0.05), but was not observed to be significantly differentially expressed in any of the wide shelf incubation time points (Fig. 10A). However, *FTN* was significantly expressed in the Fe treatment relative to the DW at T_2_ in the wide shelf (Fig. 10A). Genes associated with the nitrate assimilation response of diatoms showed an inverse relationship with indications of DFB-induced Fe limitation in both of the incubations (Fig. 10). As observed, the relative expression of *NR* and *nirA* was significantly higher in the Fe treatments than those of the DFB treatments at T_2_ for both incubations (p-adj < 0.05).

**Figure 10.**
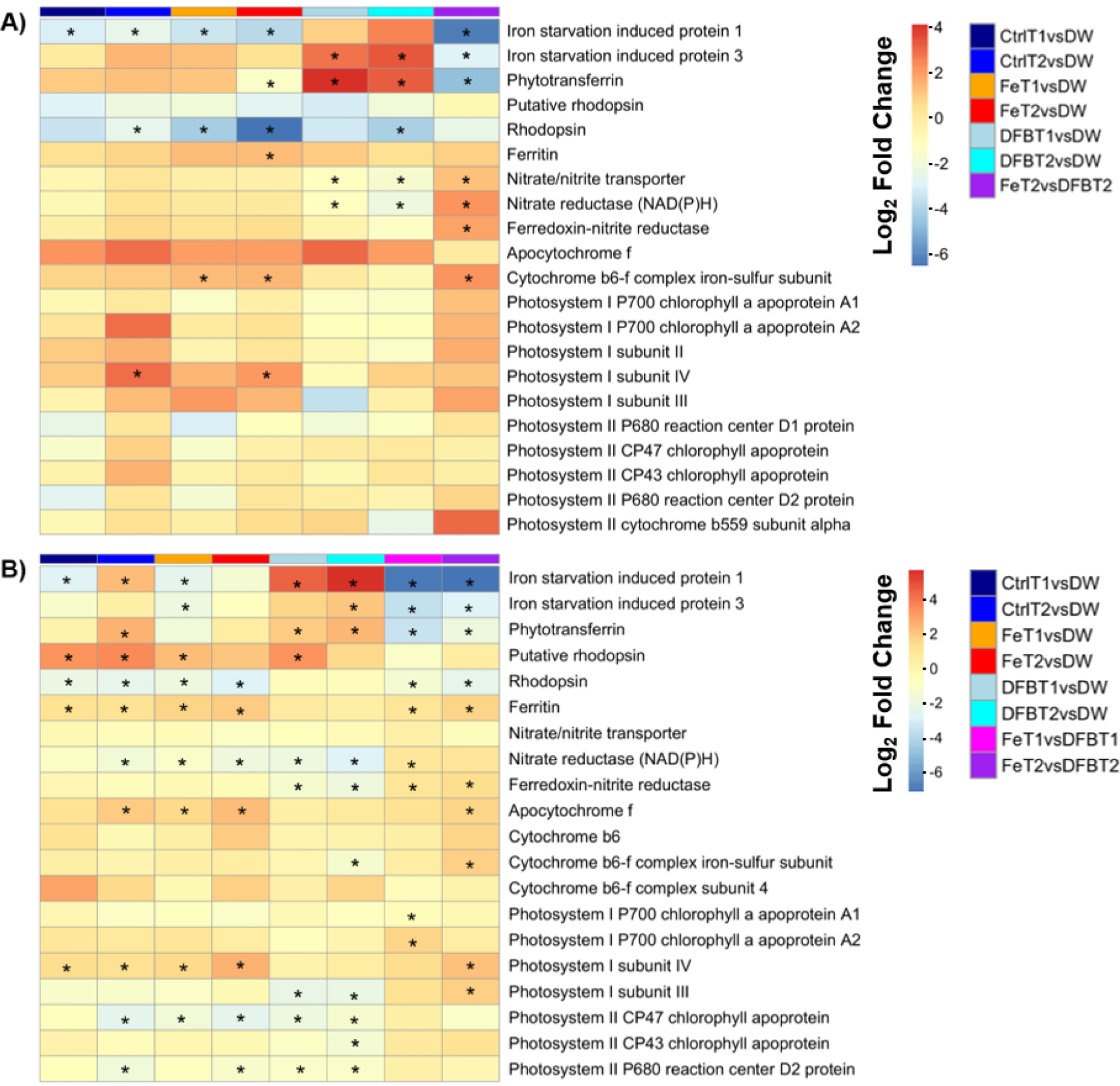
Log2 fold change of select Fe starvation, Fe storage, nitrate assimilation and photosynthetic genes between various treatment groups with respect to DW, T1, and T2 of A) wide shelf diatoms and B) narrow shelf diatoms. Significantly differentially expressed genes (log2 fold change > 1 or < -1, p-adj < 0.05) are denoted by an asterisk (*).

In both incubations, diatom photosynthesis genes were comparably highly expressed in the Fe treatments relative to the DFB treatments at T_2_. Photosystem subunit genes (*psaB, psaE, and psaF*) were all found to be significantly expressed in the Fe treatment at either time points in the narrow shelf incubation, while the cytochrome b6-f complex genes (*petA*, *petB*, *petC*) were significantly highly expressed only at T_2_ in the narrow shelf incubation. Diatom differential expression of photosynthesis genes between the Fe and DFB treatments were observed in the wide shelf incubations at T_2_, where *petC* increased in expression in the Fe treatment relative to the DFB treatment at both time points (Fig. 10A)

## Discussion

Our findings suggest a larger initial biomass of the deep-water (T_0_) community in the narrow shelf incubation likely played a role in shaping the enhanced physiological processes of the phytoplankton by T_1_ of the incubation, while the smaller initial biomass of the wide shelf contributed to a slower response to the simulated upwelling. Furthermore, neither the wide nor narrow shelves exhibited Fe limitation on a physiological level, given that physiological data between the control and Fe treatments did not significantly differ throughout the incubations. However, significant physiological and molecular differences were observed between control and DFB and Fe and DFB treatments throughout both incubations. At the molecular level, we found that diatom nitrogen assimilation and photosynthetic genes were significantly highly expressed in the Fe treatments, while Fe-starvation indicating genes were significantly highly expressed in the DFB treatments. Furthermore, the comparison of the deep-water diatom communities between the wide and narrow shelves showcased significantly increased expression of the Fe-starvation induced protein (ISIP3) indicator gene in the narrow shelf, suggesting the likelihood that Fe limitation at a molecular level may have been occurring before it was observed on a physiological level.

### Physiology and community composition changes due to simulated upwelling

Chlorophyll *a* concentrations in T_2_ of both incubations suggest that large cells (≥5 µm) comprise much of the phytoplankton biomass within the community as a result of simulated upwelling, such that this fraction was almost an order of magnitude higher than that of the small cell fraction (<5 μm). This is further substantiated by the phytoplankton taxonomic composition based on both normalized RNA and 18S rDNA read counts, where diatoms comprise a majority of the microalgal populations at T_2_ in both the wide and narrow shelf incubations despite a minor presence in the respective initial deep-water communities. This is consistent with the concept that diatoms are known to dominate phytoplankton communities in coastal upwelling regions [Armbrust 2009, Malviya et al. 2016]. Additionally, similarities in chlorophyll *a* concentrations and DIC and NO_3_ uptake rates in the control and Fe treatment indicate that neither communities in the wide shelf or the narrow shelf were experiencing Fe limitation on a physiological level in the initial stages of the simulated upwelling-induced phytoplankton blooms. Our Fe addition results corroborate various Fe fertilization experiments performed previously [Aumont and Bopp 2006, Coale et al. 1996]. The DFB (Deferroxamine B) treatments, in which a strong organic chelator (i.e., siderophore) that binds to Fe was added [Kiss and Farkas 1998], were used to decrease Fe bioavailability. Interestingly, the dissolved Fe concentration measured from the same incubations was higher in concentration in the presence of the siderophore (Supplemental Fig. 4). This is likely because DFB enhances Fe solubility and mediates the transfer of Fe from the particulate to the soluble pool (Segovia et al. 2017). However, Fe speciation plays an important role in Fe bioavailability in addition to dissolved Fe concentrations, and this Fe in the DFB treatments is likely not bioavailable because it is bound to the strong chelator. The DFB treatment resulted in expectedly lower phytoplankton growth rates and biomass accumulation relative to the control and Fe treatments in both incubations. Likewise, the addition of Fe or DFB differentially affected the physiology and growth kinetics of the larger phytoplankton, most evidently at the second timepoint. However, these size-fractionated physiological measurements only provide a coarse overview of differential responses among two phytoplankton size classes; future applications should employ innovative strategies such as pairing fluorescence-assisted cell sorting (FACS) with stable isotope analysis to provide for a higher taxonomic resolution to our understanding of rate processes among various phytoplankton functional groups. Previous studies [Casey et al. 2007, Fawcett et al. 2011] have shown the feasibility of coupling FACS with stable isotope analysis to assess uptake kinetics of phytoplankton groups discernible from flow cytometry, such as cyanobacteria and pico-eukaryotes, but expanding these applications to larger cells such as diatoms would greatly improve our knowledge of taxonomic group-based biological rate processes and nutrient consumption in these coastal upwelling zones.

Fluorescence-activated cell counts using the BD FACS Melody also suggest a direct negative effect of DFB-induced Fe limitation on the cell abundances of pico-eukaryotes, smaller diatoms, and larger centric diatoms in the wide shelf at the second time point. Notably, cells in the control did not exhibit slower growth rates than those of the Fe treatments, further supporting our observation that both communities in the wide and narrow shelves were not experiencing Fe limitation under ambient conditions. Similar studies conducted in the region have shown that Fe addition does not necessarily enhance growth relative to the natural community, but instead enhances chain forming morphology of centric diatoms [Mackey et al. 2014]. Nonetheless, both pennate and centric diatoms outperformed the rest of the phytoplankton community, as indicated by the increases to their DIC and NO_3_ uptake rates, cell concentrations, and relative transcript abundances by the second time point during both incubation experiments. Furthermore, nitrogen assimilation gene (e.g., *NRT*, *NR*, *nirA*) transcript abundances are shown to be higher in the deep-water diatom community relative to the rest of the incubations in both sites. This pattern is seldom seen in the other three major phytoplankton taxa, especially dinoflagellates, where the same nitrogen assimilation gene transcript abundances are higher towards the end of the incubations. The data presented is consistent with previous studies that show a pro-active upwelling response in diatoms while other phytoplankton taxa exhibit a reactive response to changes in environmental conditions associated with the upwelling conveyor belt cycle [Lampe et al. 2018; Lampe et al. 2021].

Within the diatoms, the smaller diatom taxa in the narrow shelf appeared to sustain population growth at T_2_ despite the DFB treatment, suggesting the possibility that the small diatoms may have been acclimated and/or buffered against low Fe conditions. This is further substantiated by the taxonomic distribution of *Pseudo-nitzschia*, which sustained their normalized RNA transcript and 18S rDNA read abundances in spite of the DFB treatment at T_2_ of the narrow shelf incubation, while *Thalassiosira* decreased its relative proportion of RNA transcripts and 18S rDNA reads at the same time point and treatment. One explanation for this phenomenon could be that pennate diatoms, especially *Pseudo-nitzschia*, which comprise the smaller diatom group, can utilize the Fe-storage protein ferritin during high Fe conditions to better buffer against episodes of low Fe and sustain growth when Fe becomes scarce [Lampe et al. 2018, Marchetti et al. 2009, Sugie et al. 2011, Yoshida et al. 2006]. Likewise, the ability of *Pseudo-nitzschia* to continue cell divisions even in the presence of low dissolved Fe availability due to the presence of the siderophore could suggest the use of such an Fe storage strategy. In contrast, many large centric diatoms that lack ferritin will be unable to sustain growth in the low dissolved Fe availability, DFB treatment [Kustka et al. 2007]. Interestingly, while deep-water communities of *Pseudo-nitzschia* maintained a relatively high abundance of ferritin gene transcripts in both the wide and narrow shelves, *Thalassiosira* displayed a lower ferritin gene transcript abundance in the deep-water of the narrow shelf. This may perhaps support the physiological evidence that *Pseudo-nitzschia* were able to sustain growth at T_2_ of the DFB treatment in the narrow shelf incubation, while *Thalassiosira* did not. Ultimately, the viability of these smaller pennate diatoms under Fe limitation may have major implications for diatom composition shifts in future Fe-limited waters. In particular, *Pseudo-nitzschia* are known to produce domoic acid [Hasle 2002], a neurotoxin that may amplify in the food web through bioaccumulation and cause harm to humans and marine mammals alike. If indeed Fe limitation induces shifts in diatom composition towards *Pseudo-nitzschia* favorable conditions, the economic and environmental damage that these toxin-producing diatoms may impose in coastal regions could compound with various other climate change consequences.

In the wide shelf incubation, biomass-normalized DIC uptake in the smaller phytoplankton size-fraction (<5 μm) did not significantly increase by the first time point but was observably higher at the second time point, while biomass-normalized nitrate uptake rates of the small cells decreased in all treatment groups by T_1_ and only marginally increased by T_2_ in the control and Fe treatments. Similar to chlorophyll *a* trends, biomass-normalized uptake rates of both DIC and NO_3_ in the large size-fraction were higher in the control and Fe treatments by T_2_ in both incubations, while DFB treatments resulted in significantly lower uptake rates. However, DFB treatments in the small size-fraction did not seem to have a significant effect on DIC or NO_3_ uptake rates during the narrow shelf incubation. This is consistent with the fact that smaller cells have a competitive advantage over larger cells for Fe uptake, as they have relatively lower demand for Fe and are able to acquire Fe reserves faster than larger cells [Finkel et al. 2010, Wells 1999]. These findings are consistent with previous studies that also suggest a potential community shift towards smaller phytoplankton size classes in future oceans, a concerning trend that may have cascading negative effects on productivity and food web stability [Finkel et al. 2010].

Comparing across the incubations, the narrow shelf incubation started with higher biomass from the initial timepoint, and by T_2_ exhibited higher phytoplankton biomass than the wide shelf incubation by approximately an order of magnitude across all treatments in the large size-fraction. Interestingly, rate processes continued in an upward trend in the wide shelf incubation as the cells were slowly acclimating to simulated upwelling; in contrast, large phytoplankton NO_3_ uptake rates in the narrow shelf incubation appeared to be shifting down by the second time point. This is also supported by the differences in the macronutrient data compared across the two sites, where the drawdown of NO_3_, PO_4_, and Si(OH)_4_ is more evident in the narrow shelf incubation by T_2_. It was difficult to discern differences in macronutrient drawdown in the wide shelf incubation because there was slower biomass accumulation as a result of a low initial seed population biomass. Likewise, the larger initial seed community within the narrow shelf was able to rapidly increase growth in preparation for the simulated upwelling. Reasons behind the large seed population density at the narrow shelf are currently unknown, but a few explanations may be considered: (1) The deep-water community may have been advected horizontally from a different location with a larger seed population. (2) The onset of upwelling-induced water column mixing prior to the incubation may have contributed to an increased vertical spatial homogeneity in cell abundances below the euphotic zone. (3) Biological differences in phytoplankton development in the narrow shelf may have provided for greater success in their acclimation to upwelling relative to the seed populations in the wide shelf. This last explanation could be summarized as an innate ability of certain diatoms to utilize the mixing and current patterns of upwelling cycles to seed upwelling systems, in which higher sinking rates have been correlated with a greater advantage to resting spore formation [Pitcher 1990]. In essence, increased sinking rates could quickly remove the nutrient depleted population from an inhospitable environment towards the end of the upwelling conveyor belt cycle, and may prevent any horizontal movement of the community so that cells are maintained at the center of the upwelling conveyor belt cycle to be primed for the next upwelling event. Further examination of gene expression of diatoms during stationary phase following an upwelling-induced diatom bloom could help decipher their ability to modulate sinking and buoyancy.

Additionally, a slight “shift-down” effect in nitrogen assimilation processes at the second time point in the control treatment during the narrow shelf incubations would indicate the possibility that the community was starting to experience physiological stress, potentially due to low Fe concentrations. Indeed, the macronutrient concentrations as well as the subsequent drawdown in the narrow shelf region had striking similarities to that of previous studies [Bruland et al. 2001], where a range of 16 – 18 μM of initial NO_3_ concentrations decreased by only a small amount. Our incubations at the narrow shelf seemed to have drawn down only slightly more NO_3_ in the control (4.4 μmol L^-1^ drawdown) and Fe (5.3 μmol L^-1^ drawdown) treatments.

### Diatom gene expression analysis

By examining diatoms at the molecular level, we found significant differential expression of genes between control and DFB and Fe and DFB treatments using Principal Component Analysis (PCA). Diatoms are known to produce resting spores as part of their life history, and these spores act as viable seed banks for future resuspension from the sediments [McQuoid and Hobson 1996]. Conceivably, the gene expression within these seed populations are likely to be considerably different below the euphotic zone compared to the gene expression after upwelling. As nutrient uptake and photosynthesis rates are enhanced in the euphotic zone, expression of various assimilation and photosystem genes are likely to play an appreciable role in the phytoplankton upwelling response. Likewise, PCA of the samples show that control and Fe treatments clustered closer together than the DFB treatment respective of the time points, but DW genes showed the furthest distance from any time point (Supplemental Fig. 7). Certain outliers (T_2_ control triplicate in wide shelf and T_1_ DFB triplicate in narrow shelf) that did not cluster with their respective sample could be due to biological variation during the incubation process. Nonetheless, the general distinctive trends between the control/Fe clusters and the DFB clusters suggest differential responses to Fe limitation in diatoms.

As inferred through both the physiology and gene expression data, Fe addition had a net positive effect on diatom growth compared to DFB addition. The significantly increased expression of nitrogen assimilation genes (i.e., *NR*, *nirA*) and photosynthetic genes (i.e., *psaB*, *psaE*, *psaF*, *petA*, *petB*, *petC*) in the Fe treatments substantiates the evidence that Fe-addition stimulates growth in phytoplankton through enhanced nitrogen assimilation and photosynthetic activity [Geider and Roche 1994]. Sensibly, Fe limitation directly impacts diatom photosynthetic efficiencies [Greene et al. 1991, Marchetti and Harrison 1997] and N assimilation rates [Timmermans et al. 1994], with notable reductions in both aspects of their growth. The increased expression of *ISIP1*, *pTF* (*ISIP2A*), *ISIP3*, and *RHO* genes in the DFB treatments of both incubations also corresponds to the noticeable decline in DIC and NO_3_ uptake rates and biomass due to Fe limitation. These results are consistent with previous studies that have also observed increased *ISIP* and *RHO* expression in Fe-stressed diatom cells [Cohen et al. 2017]. As discussed, Fe starvation induced proteins such as *ISIP1*, *ISIP2A*, and *ISIP3* are uniquely employed by certain diatoms to cope with Fe stress [Behnke and LaRoche 2020], while the significantly increased expression of *RHO* with respect to the DFB treatment could suggest the potential usage of an Fe-independent energy production/conservation strategy by diatoms under Fe limiting conditions [Cohen et al. 2017, Marchetti et al. 2015]. Although DFB is not necessarily a quantitative predictor of future Fe limitation, it does present a scenario of how declining Fe bioavailability can influence diatom physiology and gene expression. The natural siderophore is part of a vast quantity of organic ligands in the oceans that play vital roles in controlling trace metal bioavailability, and has been shown to negatively influence phytoplankton growth rates when present at high concentrations [Sanchez et al. 2018]. Furthermore, the relatively higher number of significantly differentially expressed diatom KEGG orthologs (KO) by T_2_ of both incubations compared to the other three taxa (Haptophytes, Chlorophytes, and Dinophytes) would suggest that diatoms are more transcriptionally sensitive to shifts in Fe bioavailability (Supplemental Fig. 6). On one end, their ability to outcompete other phytoplankton for Fe [Boyd et al. 2007] has enabled them to dominate many areas of the world’s oceans where Fe is available in transient supply; on the other end, they are equipped with molecular machinery used to efficiently assimilate Fe when Fe becomes limiting. It is thus not surprising that the Fe concentrations in diatoms, as well as that of other metal quotas (e.g., Mn, Ni, Zn) relative to phosphorus, are higher than most other phytoplankton taxa [Twining et al. 2011].

Comparing across the two incubation sites, there is a clear distinction in the timing of nitrogen assimilation activity between the wide and narrow shelves. Whereas the wide shelf diatoms differentially expressed nitrogen assimilation and photosynthetic genes at T_2_, the narrow shelf diatoms were able to differentially express nitrogen assimilation and photosynthetic genes at both T_1_ and T_2_, suggesting that the narrow shelf diatoms had already exhibited a response to Fe variability within 48 hours, while the wide shelf diatoms were still acclimating to the simulated upwelling at the same time point. The significant differential expression of diatom ferritin at both time points of the narrow shelf is not evident in the wide shelf incubation at either time points, and may be explained by the fact that ferritin is associated with the luxury uptake of Fe, and thus has more consistent higher transcript abundance after cells have already undergone other photosynthetic metabolic processes [Marchetti et al. 2009]. Therefore, it may be assumed that diatoms in the wide shelf were still building photosynthetic machinery through nitrogen assimilation, as indicated by their expression of *ATPase*, *NR*, and *nirA* at T_2_ in the wide shelf incubation relative to the narrow shelf incubation (Supplemental Fig. 8). This would further reinforce our hypothesis that the larger seed population of diatoms in the narrow shelf was able to rapidly increase their growth during the simulated upwelling, while the smaller initial population in the wide shelf took longer to acclimate and thus was still in a balanced growth phase during the second time point. The differential expression of various Fe stress genes in the initial diatom communities of the wide and narrow shelves (Supplemental Fig. 8) raises another question of what specifically is driving these molecular differences across the communities. One possibility may be that the narrow shelf seed community was advected through a horizontal undercurrent and contributed to the initial larger biomass. If that is the case, the previous community that was advected may have been undergoing a period of Fe limitation post bloom before reaching the area of collection, leading to the significantly increased expression of *ISIP3* and *psaE*. More attention on time histories of water masses could help in our understanding of the differences in the physiological and molecular dynamics of these seed communities between the wide and narrow continental shelves.

Consistent with the indication from the physiological data that there seemed to be a “shift-down” response in the control and Fe treatment at T_2_ in the narrow shelf, gene expression trends in diatoms are consistent with Fe-limited growth, as both *ISIP1* and *ISIP3* genes were significantly differentially expressed in both the control and Fe treatments at T_2_ relative to those in the wide shelf incubation; this would likely suggest that Fe became limiting by the second timepoint of the narrow shelf incubation, and thus could have played a role in the reduced nitrate assimilation rates. Interestingly, comparisons between the deep-water communities of either shelves exhibit increased gene expression of rhodopsin (*RHO1*, *RHO2*) in the wide shelf relative to the narrow shelf, while *ISIP3* was more significantly expressed in the narrow shelf diatom seed populations relative to the that of the wide shelf. This differential expression could be explained by differences in diatom community composition across the two sites, but may also confirm previous evidence that the narrow shelf is more likely to be Fe-limited or enter a state of Fe limitation following the upwelling-induced phytoplankton bloom [Bruland et al. 2001]. If that is the case, the abundance of *Pseudo-nitzschia* in the DFB treatment at T_2_ of the narrow shelf incubation is one exception to this trend, and may possibly be explained by their consistent expression of ferritin transcripts in the deep-water population relative to the rest of the time points in the narrow shelf. Ultimately, *Pseudo-nitzschia* cells ability to sustain growth under Fe limitation may be attributed to the combined effects of the Fe-storage protein and the relatively lower demand and higher acquisition potential for Fe. Further experimentation and analysis should focus on examining transcriptional differences among the various diatom species that dominate these upwelling blooms, and verifying whether ferritin has a significant effect on *Pseudo-nitzschia* success relative to the other diatoms.

## Conclusions

Nutrient uptake rates, community composition and gene expression data have been combined to show a differential response by phytoplankton to the upwelling conveyor belt cycle across the Fe limitation mosaic of the California Current System. This topographical and biological variability within the CCS has affected a complex set of physical and biological interactions which shape the phytoplankton community structure and behavior in response to environmental changes. To address our central questions, we found that (1) Fe limitation significantly influences not only diatom bloom formation but also phytoplankton physiology, community structure, and gene expression. Fe-limited phytoplankton, especially larger sized cells, exhibited slower growth rates and carbon and nitrogen-specific uptake rates under Fe-limitation. More specifically, (2) diatom gene expression showed differential patterns in response to Fe variability, where *ISIP* gene expression increased in DFB treatments, and *FTN* gene expression increased in Fe treatments. Furthermore, (3) seed populations play an important role in shaping the phytoplankton upwelling response and acclimation time. As observed, a phytoplankton seed population with larger biomass in the narrow shelf incubation resulted in a rapid biomass accumulation rate relative to that of the wide shelf incubation. The causes for this high initial biomass may be attributed to a variety of factors, ranging from physical forces such as horizontal advection and undercurrents to biological attributes of diatoms that may adjust their sinking buoyancy. Lastly, (4) the apparent ability to sustain growth by smaller phytoplankton – including smaller diatoms – under Fe limitation may cause drastic shifts in phytoplankton composition within coastal upwelling systems. These changes in community structure could have major implications for not only the food web but also global biogeochemical cycles.

Ultimately, this study was conducted with the aim to better understand complex upwelling systems. Variations in the physical, chemical, and biological parameters of the CCS play a role in shaping how we interpreted our results. For instance, the lower biomass in the initial wide shelf community not only reduced the sample size of certain treatments but also compromised our confidence in studying the biology of their response. Nonetheless, environmental stochasticity is a fundamental aspect of ecological research, and perhaps even beneficial to our understanding of how biological processes occur in natural phytoplankton communities. The presence of climate change, however, could drastically alter phytoplankton assemblages in future oceans [Benedetti et al. 2021], and consequentially influence the composition of their seed populations. Hence, it is imperative for us to examine in further detail how environmental conditions can impact these dominant phytoplankton taxa, especially within an intricate and complex system such as the CCS, where future ocean change is projected to intensify the Fe limitation gradient between wide and narrow continental shelves.

## Acknowledgements

We are grateful for the crew and staff of the R/V Oceanus and their valuable assistance and support throughout the cruise. We also thank Dr. Ryan Paerl at North Carolina State University for providing us the BD FACS Melody Flow Cytometer to process our samples, and Brian Dreyer at University of California Santa Cruz for assistance with the ICP-MS analyses.

Funding for the project was provided to A.M. from a National Science Foundation grant (OCE1751805). The trace metal analyses were supported with startup funds from California State Polytechnic University, Humboldt and Award #26844 from Research Corporation for Science Advancement’s Cottrell Scholar Award to C.P.T.

