## Supporting Information for "Variability in the phytoplankton response to upwelling across an iron limitation mosaic within the California Current System"

**1. Materials and Methods**

*1.1 Isotope uptake of dissolved inorganic carbon and nitrate*

For the wide shelf incubation, samples were spiked with 200 μmol L^-1^ of NaH¹³CO₃ and 1 μmol L^-1^ of Na¹⁵NO₃ according to estimated ambient DIC concentrations of 2000 μmol L^-1^ and measured surface nitrate concentrations of 10 μmol L^-1^. For the narrow shelf incubation, samples were spiked with 200 μmol L^-1^ of NaH^13^CO_3_ for all samples and 1.5 μmol L^-1^ of Na^15^NO_3_ for the deep-water triplicate samples. This amount was amended according to estimated ambient DIC concentrations of 2000 μmol L^-1^ and measured nitrate concentrations of 15 μmol L^-1^ in the deep-water.

Absolute uptake rates of dissolved inorganic carbon and nitrate (ρ, DIC or NO_3_ taken up per unit time) were derived from the constant transport model (Eq. 3 from Dugdale and Wilkerson 1986). The ^13^C fraction was first calculated by dividing the measured ^13^C atom percentage in each sample by the natural ^13^C atom percentage (1.087%), over the difference between the percentage of ^13^C over total C in the sample and the natural ^13^C atom percentage. The ^13^C biomass was then derived by multiplying the ^13^C fraction by the calculated POC concentration for each sample, and ^13^C absolute uptake rate was measured as the ^13^C biomass over the incubation time (6 hours). The same calculations were applied to assess the ^15^N absolute uptake rates, but with the respective natural atom percentage of ^15^N (0.367%).

*1.2 Dissolved Fe*

Samples for dissolved Fe were taken from each cubitainer within the bubble by filtering through pre-cleaned 0.2 μm pore size polyethersulfone membrane Acropak-200® capsule filters into LDPE bottles that had been rigorously cleaned as in the GEOTRACES cookbook (Cutter et al., 2014). Sample bottles were rinsed three times with sample before filling. Samples were acidified at sea to pH ~1.7 with optima HCl (2 mL of 12 M HCl per liter of seawater) and were analyzed after the cruise with the methods of Biller and Bruland (2012). Briefly, this method involves pre-concentration onto Nobias-chelate PA1 resin followed by analysis with a High Resolution Inductively Coupled Plasma Mass Spectrometer. For quality control, a few samples were rerun with a flow injection analysis method (Lohan et al., 2006, with modifications as described in Biller et al. 2013). This method involves pre-concentration on toyopearl resin followed by in-line spectrophotometric analysis.

*1.3 DNA collection and 18S rDNA analysis*

Approximately 1L of seawater from each cubitainer was collected onto a 0.45 μm Supor-450 Pal filters (47 mm) using a vacuum pump, then immediately flash frozen in liquid nitrogen and stored at -80 C until DNA extraction in the laboratory. Prior to DNA extraction, 2.5 x 10^7^ copies of a V4 plasmid were added to each sample as an internal standard [Lin et al. 2019]. DNA was extracted using the Qiagen DNeasy Plant Mini Kit. Polymerase Chain Reaction (PCR) was performed at 98 °C for 5 min, followed 30 three-step cycles of 94 °C for 30s, 57 °C for 45 s, 72 °C for 1 min, and finally 72 °C for 2 min. PCR triplicates were pooled (20 uL from each triplicate) and purified using Qiagen QIAquick DNA Purification Kit. DNA concentration was measured using the Invitrogen Qubit 4 fluorometer and Qubit dsDNA HS Assay Kit.

Each pooled sample was then normalized to 17 nM and pooled in equal volumes to produce a compiled library sample, which was sequenced by GENEWIZ using Illumina Miseq. Reads (2x300 base pairs) were demultiplexed in QIIME2 [Bolyen et al. 2019] and Python, trimmed for quality control, and merged using the cutadapt plugin [Martin 2011]. Amplicon Sequence Variants (ASVs) were assigned using the Dada2 plug-in (QIIME2) and subsequently identified and clustered using a 97% similarity threshold [Callahan et al. 2016]. The ASVs were then taxonomically classified via the PR2 18S v.4.13.0 gene taxonomic reference dataset [Guillou et al. 2012].

**2. Results and Discussion**

*2.1 Absolute DIC and Nitrate Uptake Rates*

Absolute DIC (3.72 ± 1.98 nmol L^-1^ hr^-1^, average ± standard deviation) and nitrate (1.03 ± 0.29 nmol L^-1^ hr^-1^) uptake of the large cell (≥5 µm) deep-water community in the wide shelf incubations were lower than the absolute DIC (12.71 ± 2.781 nmol L^-1^ hr^-1^) and nitrate (2.27 ± 0.38 nmol L^-1^ hr^-1^) uptake rates of deep-water large cells in the narrow shelf incubations. For deep-water (T_0_) phytoplankton <5 µm, absolute DIC uptake (Supplemental Fig. 3B) in the wide shelf incubation (2.17 ± 0.62 nmol L^-1^ hr^-1^) was lower than that of the narrow shelf incubation (8.50 ± 4.06 nmol L^-1^ hr^-1^). Absolute nitrate uptake rates for both the wide (1.46 ± 0.91 nmol L^-1^ hour^-1^) and narrow (1.47 ± 0.54 nmol L^-1^ hr^-1^) shelf incubations were comparable in the small cell (<5 µm) deep-water community. In the wide shelf incubation, absolute DIC uptake rates of the large phytoplankton (≥5 µm) showed accelerated growth by T_2_, specifically in those of the control (182.1 ± 23.3 nmol L^-1^ hr^-1^) and Fe treatment (177.4 ± 18.4 nmol L^-1^ hr^-1^), while the DFB treatment at T_2_ exhibited considerably lower absolute DIC uptake (0.86 ± 0.04 nmol L^-1^ hr^-1^). Cells in the smaller fraction (<5 µm) also exhibited the same patterns in the control (8.25 ± 1.40 nmol L^-1^ hr^-1^) and Fe treatment (5.05 ± 2.14 nmol L^-1^ hr^-1^) with respect to the DFB treatment (1.10 ± 0.94 nmol L^-1^ hr^-1^) by T_2_, but were considerably less efficient in absolute DIC uptake than the larger phytoplankton (Supplemental Fig. 3B). Additionally, in the wide shelf incubation, absolute nitrate uptake rates of the large phytoplankton in the control (14.3 ± 0.91 nmol L^-1^ hr^-1^) and Fe treatment (14.1 ± 1.36 nmol L^-1^ hr^-1^) were significantly greater than that of the DFB treatment (1.34 ± 0.12 nmol L^-1^ hr^-1^) by T_2_ (Fig. 3B). In the wide shelf small size fractions, the DFB effect in absolute nitrate uptake between the control (1.81 ± 0.25 nmol L^-1^ hr^-1^), Fe (1.76 ± 0.43 nmol L^-1^ hr^-1^), and DFB (0.91 ± 0.36 nmol L^-1^ hr^-1^) were less nuanced but still noticeable at T_2_ (Supplemental Fig. 3D).

In the narrow shelf incubation. absolute DIC uptake rates of large phytoplankton at T_1_ already exhibited differential responses to treatment variables, where the control (278.5 ± 82.68 nmol L^-1^ hr^-1^) and Fe treatment (350.8 ± 88.97 nmol L^-1^ hr^-1^) were visibly higher in DIC uptake than that of the DFB treatment (114.7 ± 19.23 nmol L^-1^ hr^-1^). By T_2_, the absolute DIC uptake rates (Supplemental Fig. 3A) had become significantly different among the control (3317.8 ± 1006.7 nmol L^-1^ hr^-1^) and Fe (4739.8 ± 1381.1 nmol L^-1^ hr^-1^) treatment, and the DFB treatment (533.4 ± 216.8 nmol L^-1^ hr ^-1^). Cells in the small size-fraction did not seem to show significant absolute DIC uptake rate differences relative to the treatment variables in either time points of the narrow shelf incubation. Control (34.2 ± 5.22 nmol L^-1^ hr^-1^), Fe (33.0 ± 11.0 nmol L^-1^ hr ^-1^), and DFB (29.1 ± 11.0 nmol L^-1^ hr ^-1^) treatments at T_1_ and T_2_ (147.7 ± 44.27 nmol L^-1^ hr ^-1^, 166.3 ± 11.82 nmol L^-1^ hr ^-1^, 241.3 ± 87.5 nmol L^-1^ hr ^-1^, control, Fe, DFB, respectively) all exhibited similar absolute DIC uptake rates. Large phytoplankton absolute nitrate uptake rates in the narrow shelf incubation followed a similar trend as that of the DIC uptake, where the control (33.2 ± 12.7 nmol L^-1^ hr ^-1^ at T_1_; 179.8 ± 76.6 nmol L^-1^ hr^-1^ at T_2_) and Fe treatment (44.98 ± 10.59 nmol L^-1^ hr^-1^ at T_1_; 262.7 ± 137.3 nmol L^-1^ hr^-1^ at T_2_) showed enhanced nitrate uptake rate relative to that of the DFB treatment (13.1 ± 2.87 nmol L^-1^ hr^-1^ at T_1_; 23.6 ± 11.4 nmol L^-1^ hr^-1^ at T_2_) at both time points, with T_2_ exhibiting greater differences between the control/Fe and the DFB treatments. For the small size-fraction, no noticeable Fe or DFB effect on absolute nitrate uptake could be observed in the control (6.46 ± 1.10 nmol L^-1^ hr^-1^ at T_1_; 14.06 ± 8.11 nmol L^-1^ hr^-1^ at T2), Fe (6.87 ± 2.13 nmol L^-1^ hr^-1^ at T_1_; 12.96 ± 3.88 nmol L^-1^ hr ^-1^ at T_2_), and DFB (5.20 ± 1.75 nmol L^-1^ hr ^-1^ at T_1_; 17.3 ± 4.22 nmol L^-1^ hr ^-1^ at T_2_) at either time point (Supplemental Fig. 3D).

*2.2 Main output of the two-way ANOVAs*

Statistical analysis of phytoplankton physiology (absolute and biomass-normalized uptake rates of DIC and nitrate) based on post-hoc ANOVA Tukey tests suggest that neither shelves exhibited iron limitation. As indicated, no significant differences in uptake kinetics were found between the control and Fe treatments in either of the phytoplankton size fractions throughout the incubation, but significant differences (p < 0.05) are observed between the control and DFB treatments as well as the Fe and DFB treatments in the large cells (≥5 μm) of both shelves (Supplemental Table 1). For the small cells (<5 μm), the only significant difference in physiology was found in the normalized DIC uptake of Fe versus DFB treatments (p = 0.029), while no other instances of significant difference between the treatments are observed. Macronutrient concentration differences across the incubation time points were also found to be statistically insignificant with respect to the control, Fe, and DFB treatments (Supplemental Table 1), supporting the interpretation that increases in macronutrient concentrations throughout the incubation may be attributed to variability in the cubitainers as a result of low biomass accumulation.

*2.3 Fe concentrations*

Dissolved Fe (dFe) concentrations in both incubations were respectively lower in the control relative to the Fe and DFB treatments (supplemental Fig. 4). In the wide shelf incubation, no observable difference in dFe concentration can be found between Fe and DFB treatments. In the narrow shelf incubation, the DFB treatments resulted in the highest concentrations of dFe by T_2_ relative to the control and Fe treatments. This is inversely reflected in the particulate Fe (pFe) concentrations at T_2_, where pFe concentrations in the DFB treatments were much lower than the control and Fe treatments. The increase of dFe concentrations in the presence of DFB maybe attributed to the likelihood that DFB enhances Fe solubility, and mediates the transfer of Fe from the particulate to the soluble pool (Segovia et al. 2017).

*2.4 Taxonomic Composition based on 18S ribosomal DNA*

A large dinoflagellate composition (79%) can be seen among the initial seed community in the wide shelf incubation. While diatoms in this initial community comprised of a small proportion (2%), they eventually made up a majority of the phytoplankton population by 120 hours (T_2_) of the wide shelf incubation in both the control and Fe treatments, while DFB exhibited a negative effect on their overall representation at T_2_ (Supplemental Fig. 5A). Diatoms in the initial seed community of the narrow shelf incubation were comparatively more abundant based on 18S rDNA reads (30%) while dinoflagellates still encompassed a small majority of the entire phytoplankton community (38%). However, a relatively quicker response to simulated upwelling can be observed at just 48 hours (T_1_) of the incubation, where diatoms are seen to have dominated the bloom in the control, Fe, and DFB treatments. Interestingly, DFB did not seem to exert a negative effect on diatom proportions in the narrow shelf incubation, in contrast to our RNA transcript data (Supplementary Fig. 5A)

Within the diatom taxa, 18S rDNA counts show that *Thalassiosira* species again represented the largest diatom proportions throughout the wide shelf incubation, with DFB having a slightly negative effect on their representation at T_2_. Contrastingly, *Pseudo-nitzschia* make up a relatively larger proportion of the initial diatom seed community in the narrow shelf incubations when compared to the two centric diatoms. Additionally, while DFB reduced the *Thalassiosira* proportions by T_2_, *Pseudo-nitzschia* proportions seem to be relatively unaffected under induced iron limitation (Supplemental Fig. 5B).

*2.5 KEGG Ortholog comparisons between wide and narrow shelf incubations*

Comparing across the incubation sites, the number of significant differentially expressed KEGG Orthologs (KO) between Fe and DFB treatments (log_2_ fold change >1 or <-1, p-adj < 0.05) were observably lower at T_1_ in the wide shelf compared to the same time point in the narrow shelf incubation, especially in diatoms (8 in wide shelf, 63 in narrow shelf), chlorophytes (1 in wide shelf, 15 in narrow shelf), and haptophytes (0 in wide shelf, 41 in narrow shelf) (Fig. 8). By T_2_ of the wide shelf incubation, the number of significant differentially expressed KOs increase drastically for diatoms (116), but was not the case for haptophytes and chlorophytes. Significant differentially expressed KOs in dinoflagellates increased only marginally from T_1_ (1) to T_2_ (13). In the narrow shelf incubation, the number of significant differentially expressed KOs are high among diatoms (140), haptophytes (96), and chlorophytes (134) at T_2_; dinoflagellates also increased in the number of these KOs from T_1_ (3) to T_2_ (28). Furthermore, a number of the same KOs are shared and conserved amongst the different taxa at T_2_ in the narrow shelf (Supplemental Fig. 6).

*2.6 Principal Component Analysis*

In both incubations, diatom genes differentially expressed in the initial deep-water community are vastly different from any of the respective treatments in the later time points (Supplemental Fig. 7). In the narrow shelf incubation, triplicate samples are observed to cluster with respect to their Fe status and time point, where the largest variance is detected amongst control, Fe, and DFB treatments at T_2_. The single data points in the wide shelf for T1 (control, Fe, and DFB) and T2 (DFB) were a result of low RNA yields, and thus were pooled into singular samples.

*2.7 Comparison between diatom deep-water seed communities of wide and narrow shelf incubations*

The diatom deep-water seed communities seemed to differ quite significantly in gene expression patterns within the initial communities prior to simulated upwelling incubations. The seed populations in the wide shelf had higher normalized transcript abundance for *RHO* with respect to the narrow shelf, as well as the gene for DNA repair protein *RAD50* (p-adj < 0.05) (Supplemental Fig. 8). In the narrow shelf, the diatom seed populations exhibited higher transcript abundance of *ISIP3* and *psaE* genes compared to their wide shelf counter parts, as well as cob (I) alamin adenosyltransferase (*MMAB*, *pduO*, p-adj < 0.05). Additionally, differential expression compared between T_2_ of either incubation suggests a higher expressionof *ISIP1* and *ISIP3* genes in the control and Fe treatments of the narrow shelf incubation relative to the wide shelf incubation (p-adj < 0.05). In contrast, diatoms at T_2_ of the wide shelf incubations exhibited significantly higher expression of ATPases (*ATPeF0F*, *ATP5J2*), *NR*, and *nirA* (p-adj < 0.05).

**Supplemental Table 1.** Statistical analysis based on post-hoc ANOVA Tukey tests of A) wide shelf and B) narrow shelf incubations. Three comparisons (control vs. Fe, control vs. DFB, Fe vs. DFB) were made between DIC and nitrate uptake (absolute and normalized) rates in either large (≥5 µm) or small (<5 µm) cell size-fractions collected from the incubation. Comparisons were also made across both treatments and time points for C) macronutrient concentrations in the wide shelf incubation.

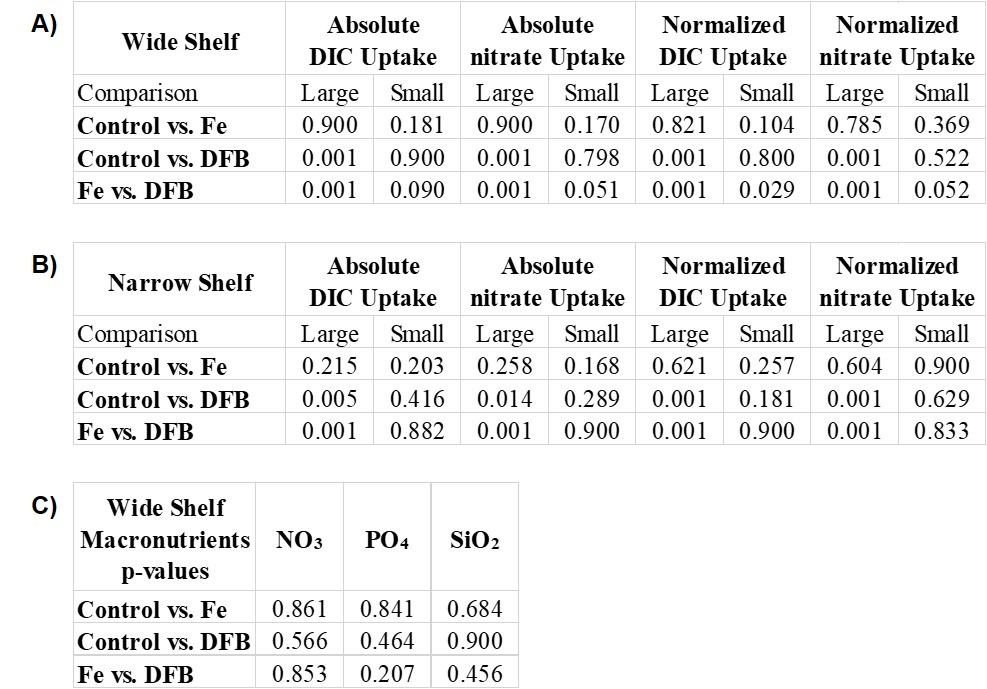

**Supplemental Table 2**. A) chlorophyll *a* averages (μg L^-1^) of the large cell (≥5 μm) and small cell (<5 μm) deep-water seed communities (T_0_) at the wide and narrow shelf incubations. B) flow cytometry-derived cell count averages (cells mL^-1^) of *Synechococcus*, pico-eukaryotes, smaller diatom, and large centric diatom, and other phytoplankton seed communities (T_0_) at the wide and narrow shelf incubations.

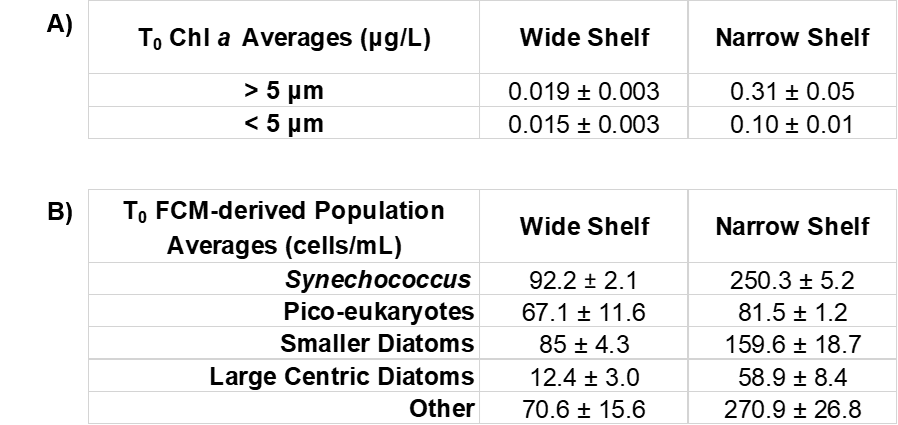

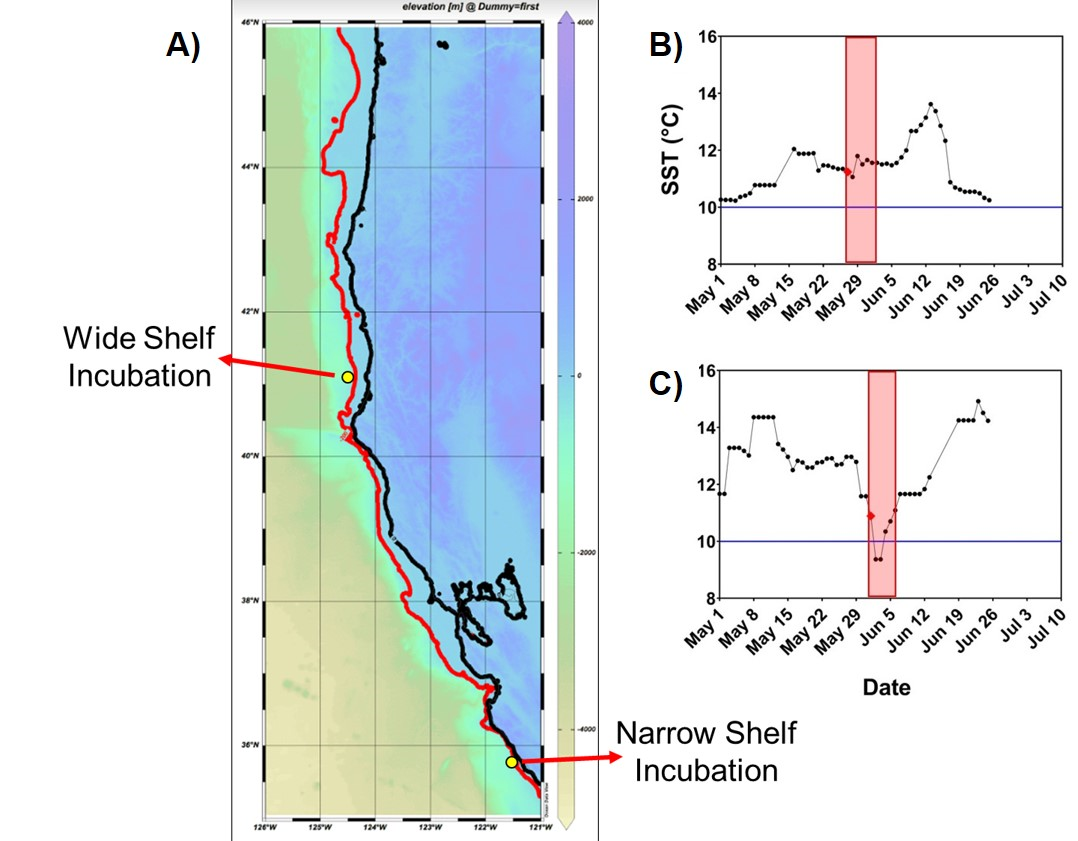

**Supplemental Figure 1**. A) Bathymetry map of the study region with the locations of the wide and narrow shelf incubations indicated by the yellow circles. The red line depicts the edge of the continental shelf. Eight-day average sea surface temperatures (°C) plotted before, during, and after deep-water collection for the B) wide shelf incubation and C) narrow shelf incubation. The first day of incubation is denoted by the red dot corresponding to May 27^th^ 2019 for the wide shelf and June 2^nd^ 2019 for the narrow shelf. The horizontal blue line is used to denote the 10 °C threshold for upwelled water, as samples were targeted to be collected during periods of relaxation. Incubation days are shaded in red. SST data after the incubation period is shown in order to provide context for the region.

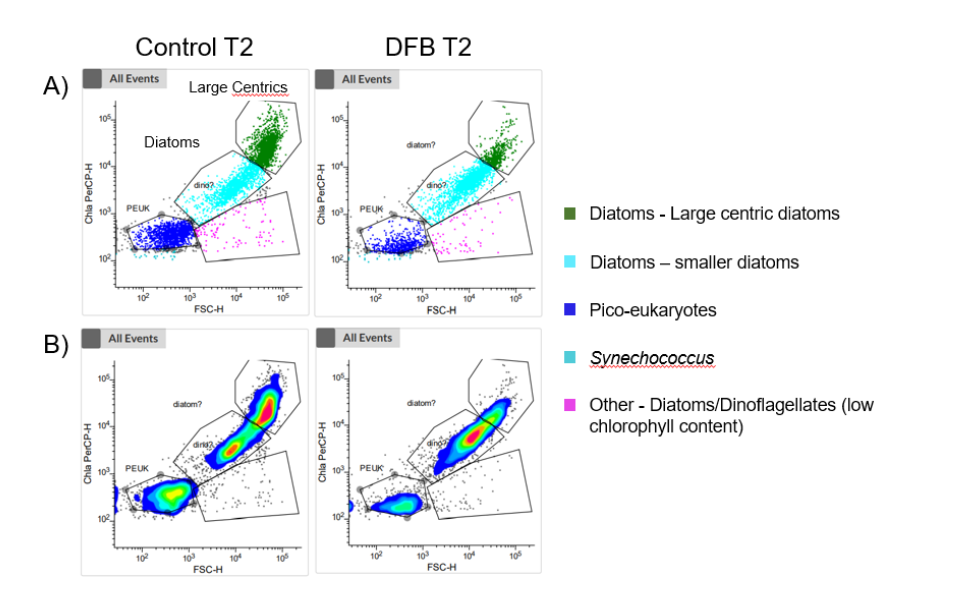

**Supplemental Figure 2**. Example scatterplots A) of one triplicate sample from the control and DFB treatments at time point 2 of the narrow shelf incubation and contour plots B) of the respective scatterplots for the same samples. Gating parameters were calibrated using a mock community containing dinoflagellates, centric and pennate diatoms, coccolithophores, and chlorophytes.

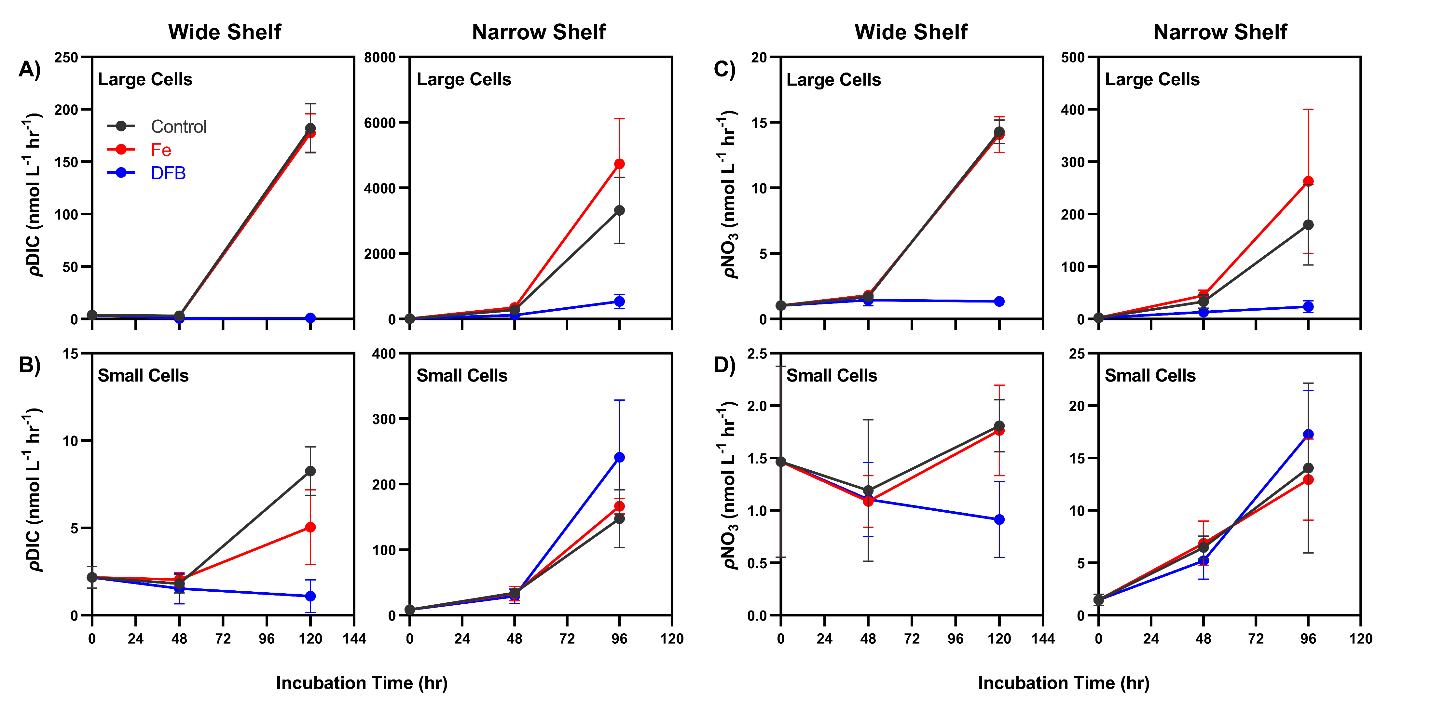

**Supplemental Figure 3**. Absolute (ρ) uptake rates of dissolved inorganic carbon (DIC) in the A) large cell size-fractions (≥5 μm) and B) small cell size-fractions (<5 μm), and absolute (ρ) uptake rates of nitrate (NO₃) in the C) large cell size-fractions and D) small cell size-fractions over the course of the incubation at the wide and narrow shelves. Grey, red and blue symbols and lines represent the control, Fe, and DFB treatments respectively. Error bars represent the standard deviations of the mean (n=3).

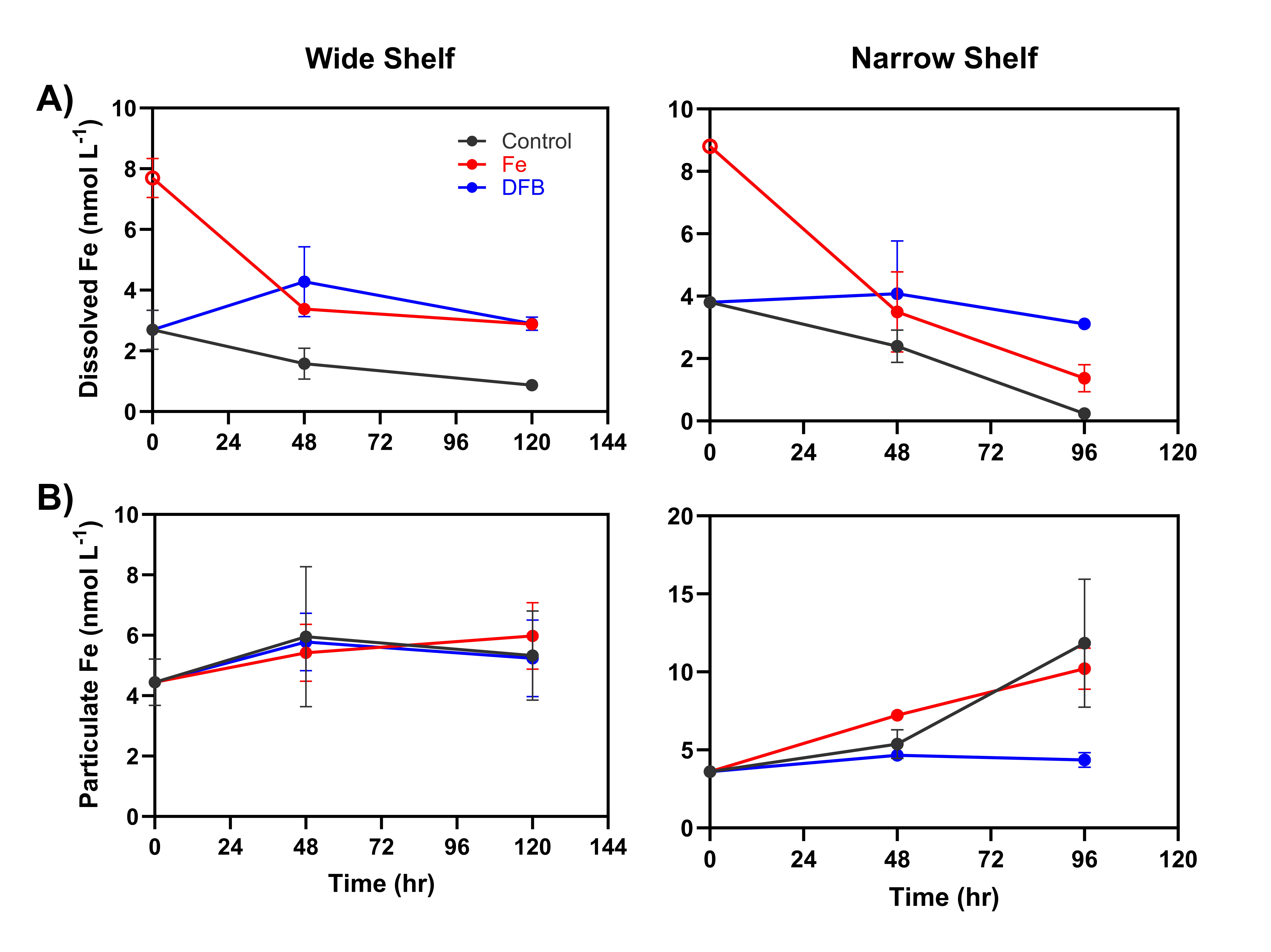

**Supplemental Figure 4**. A) Dissolved Fe concentrations and B) particulate Fe concentrations (nmol L^-1^) of the wide and narrow shelf incubations over the course of 120 and 96 incubation hours, respectively. Grey, red and blue circles and lines represent values of the control, Fe and DFB treatments, respectively. 5 nM of dissolved Fe was added to the ambient Fe concentration measured at T_0_ (red open circles) of the Fe treatment in order to estimate the total dissolved Fe concentration at this time point. Error bars represent the standard deviations of the mean (n=3).

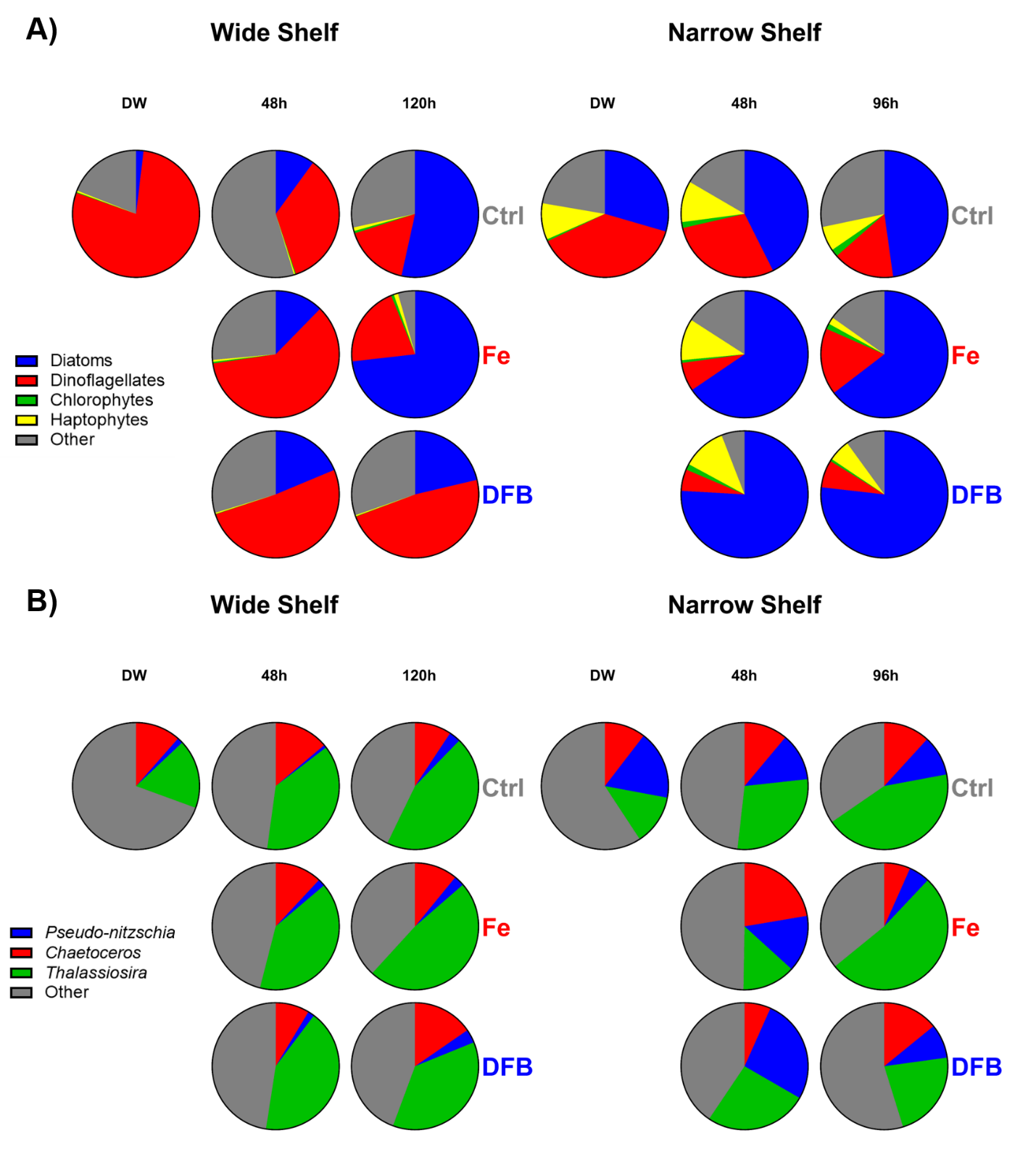

**Supplemental Figure 5**. Average taxonomic distribution by 18S ribosomal DNA (rDNA) of phytoplankton from each time point (DW: T_0_; 48h: T_1_; 120h/96h: T_2_) and treatment (Control, Fe, and DFB, respectively). Taxonomic distributions are categorized as A) percentage of mapped reads from the whole community and B) percentage of mapped reads for diatom genera within all reads assigned and subset for diatoms (Bacillariophyta).

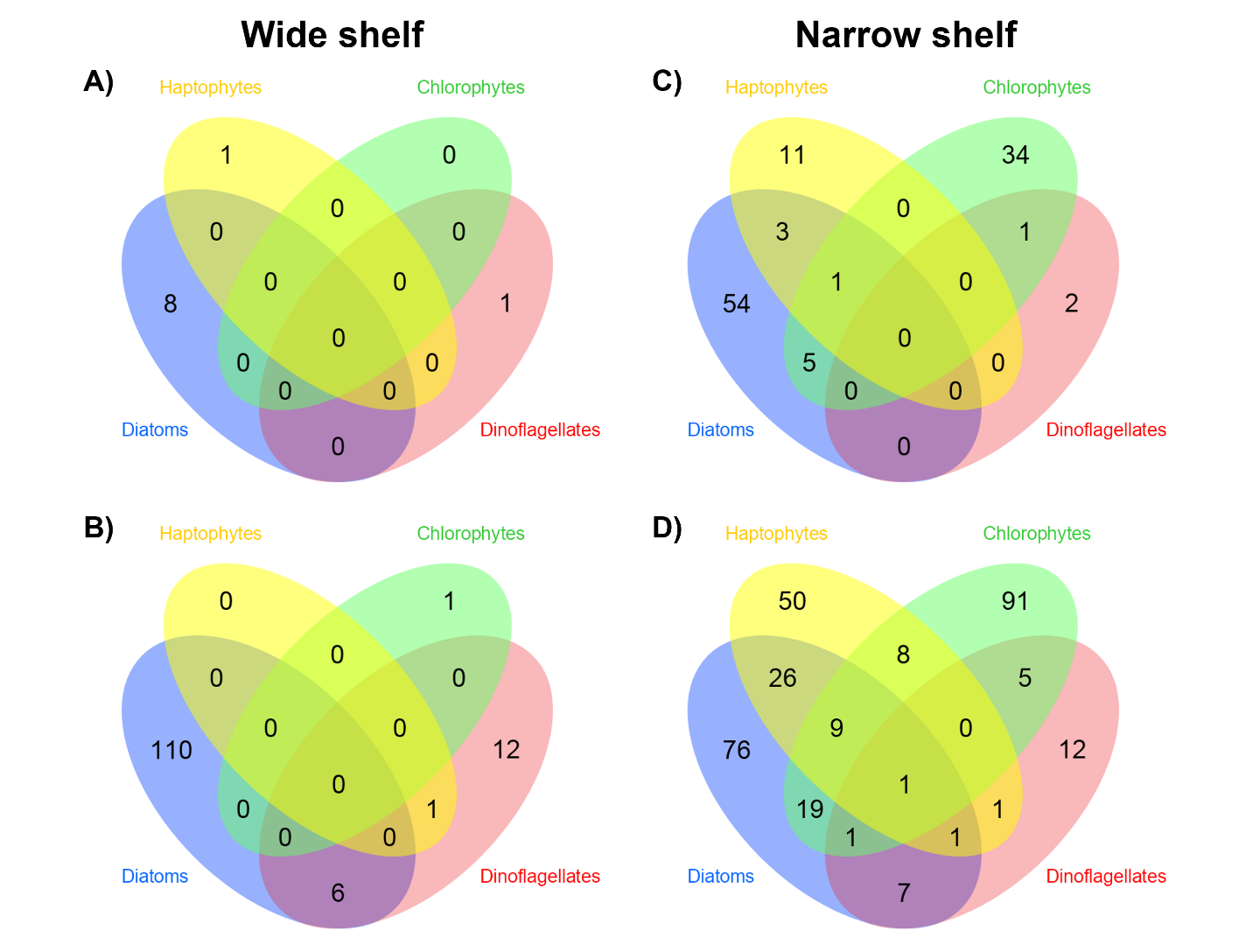

**Supplemental Figure 6**. Venn diagrams of significant (log_2_ fold change > 1 or < -1, p-adj < 0.05) differentially expressed KEGG Orthologs (KO) for the Fe and DFB treatments in the wide shelf at A) T_1_, B) T_2_, and in the narrow shelf at C) T_1_, D) T_2_ for each main phytoplankton group.

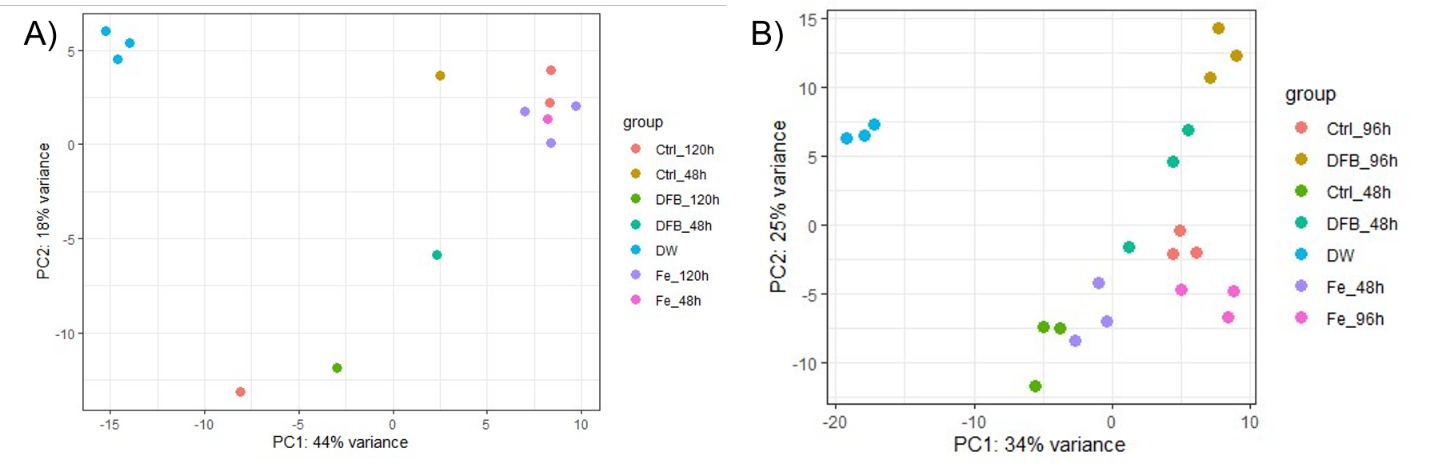

**Supplemental Figure 7**. Principle Component Analysis (PCA) plots of the top 500 most differentially expressed genes in diatoms in the A) wide shelf incubations and B) narrow shelf incubations.

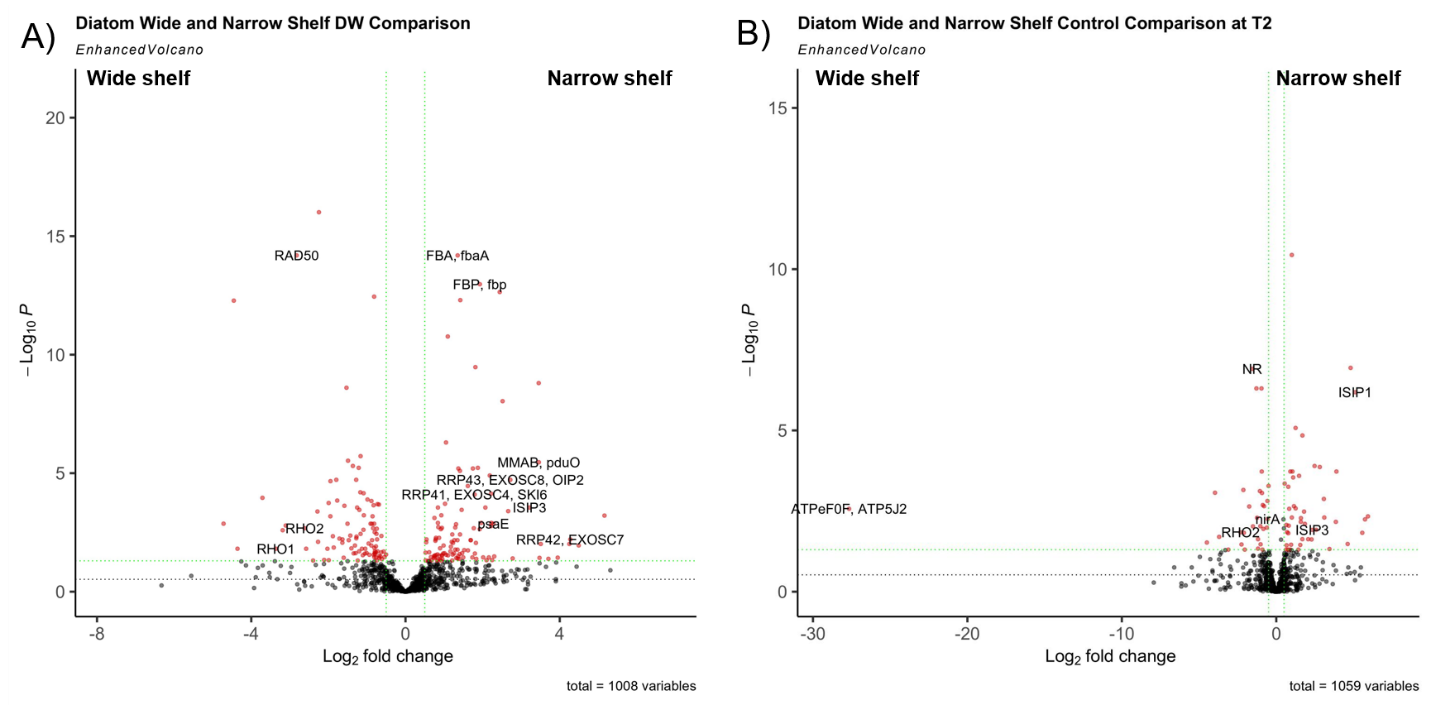

**Supplemental Figure 8**. Volcano plots of significant (log_2_ fold change > 1 or < -1, p-adj < 0.05) differentially expressed genes for A) narrow shelf seed population versus wide shelf seed population and B) narrow shelf versus wide shelf control treatments at T_2_. Genes with higher expression in the narrow shelf are shown on the right side of both plots, while genes with higher expression in the wide shelf are shown on the left side of both plots. Significance of differential expression is assessed on the y-axis, which is a logarithmic function of the adjusted p-value, such that a high significance to the differential expression would result in a higher placement on the plot.
